# Genetic population structure constrains local adaptation in sticklebacks

**DOI:** 10.1101/2020.01.17.908970

**Authors:** Petri Kemppainen, Zitong Li, Pasi Rastas, Ari Löytynoja, Bohao Fang, Jing Yang, Baocheng Guo, Takahito Shikano, Juha Merilä

**Affiliations:** Ecological Genetics Research Unit, Organismal and Evolutionary Biology Research Programme, Faculty of Biological and Environmental Sciences, FI-00014 University of Helsinki, Finland; CSIRO Agriculture & Food, GPO Box 1600, Canberra, ACT 2601, Australia; Institute of Biotechnology, FI-00014 University of Helsinki, Finland; Chinese Sturgeon Research Institute, Three Gorges Corporation, Yichang, 443100, China; The Key Laboratory of Zoological Systematics and Evolution, Institute of Zoology, Chinese Academy of Sciences, Beijing, China; Division of Ecology and Biodiversity, Kadoorie Building, The University of Hong Kong, Pokfulam Road, Hong Kong, SAR

**Keywords:** convergent evolution, epistasis, local adaptation, pelvic reduction, Pitx1, *Pungitius pungitius*

## Abstract

Repeated and independent adaptation to specific environmental conditions from standing genetic variation is common. However, if genetic variation is limited, the evolution of similar locally adapted traits may be restricted to genetically different and potentially less optimal solutions or prevented from happening altogether. Using a quantitative trait locus (QTL) mapping approach, we identified the genomic regions responsible for the repeated pelvic reduction (PR) in three crosses between nine-spined stickleback populations expressing full and reduced pelvic structures. In one cross, PR mapped to linkage group 7 (LG7) containing the gene *Pitx1*, known to control pelvic reduction also in the three-spined stickleback. In the two other crosses, PR was polygenic and attributed to ten novel QTL, of which 90% were unique to specific crosses. When screening the genomes from 27 different populations for deletions in the *Pitx1* regulatory element, these were only found in the population in which PR mapped to LG7, even though the morphological data indicated large effect QTL for PR in several other populations as well. Consistent with the available theory and simulations parameterised on empirical data, we hypothesise that the observed variability in genetic architecture of PR is due to heterogeneity in the spatial distribution of standing genetic variation caused by >2x stronger population structuring among freshwater populations and >10x stronger genetic isolation by distance in the sea in nine-spined sticklebacks as compared to three-spined sticklebacks.

## Introduction

Failure to evolve in response to changing environmental conditions may lead to extirpation or even extinction (Orr and Unckless 2008). If heritable variation necessary for adapting to environmental change already exists in the form of standing genetic variation, genetic adaptation may proceed swiftly, at least compared to the time it would take for populations to adapt from novel mutations (Orr and Unckless 2008; Barrett and Schluter 2008; Thompson *et al*. 2019). Furthermore, when exposed to novel yet similar environments, populations derived from the same ancestral population – hence carrying the same pool of alleles – can often be expected to respond to similar selection pressures in a similar fashion, leading to parallel phenotypic evolution (Arendt and Reznick 2007; Schluter and Conte 2009; Elmer and Meyer 2011; Stern 2013; Conte *et al*. 2015; Bölnick *et al*. 2018; Hermisson and Pennings 2017). However, in genetically highly structured species, potentially advantageous rare alleles may be lost due to founder events and random genetic drift, thus preventing adaptation. Alternatively, due to heterogeneity in the distribution of standing genetic variation, adaptation to given selection pressures could more likely be acquired with phylogenetically independent alleles (rather than alleles that are identical by descent) at the same or different loci influencing the same trait, even if they may differ significantly in their fitness effects (Cohan 1984; Merilä 2013, 2014; Rosenblum *et al*. 2014). Epistatic interactions and the order at which beneficial alleles enter and become established in a population also play important roles in determining the genetic architecture of the trait in question (Orr 1998). Thus, the demographic history of populations likely plays a central role in determining the likelihood of local adaptation, and hence also parallel phenotypic evolution.

In recent years, the nine-spined stickleback (*Pungitius pungitius*) has been emerging as a model system for studying adaptive evolution in the wild (Merilä 2013) alongside the more widely studied three-spined stickleback (*Gasterosteus aculeatus*; Bell and Foster 1994; Gibson 2005). The latter species’ ability to rapidly adapt to local environmental conditions has often been shown to stem from a global pool of ancestral standing genetic variation (Schluter and Conte 2009; Jones *et al*. 2012; Terekhanova *et al*. 2014, 2019). The nine-spined stickleback differs from the three-spined stickleback by having smaller effective population sizes (*N_e_*), reduced gene flow in the sea, and a tendency to occur in small landlocked ponds in complete isolation from other populations (Shikano *et al*. 2010a; DeFaveri *et al*. 2012; Merilä 2013; this study). Given their contrasting population demographic characteristics, three- and nine-spined sticklebacks can thus be expected to respond differently with respect to local adaption to newly colonised freshwater habitats.

Regressive evolution of the pelvic complex has occurred in freshwater populations of three (*viz., Gasterosteus, Pungitius*, and *Culaea*) of the five recognised stickleback genera since the last glacial period (reviewed in Klepaker *et al*. 2013). While marine populations of three- and nine-spined sticklebacks usually have complete pelvic structures with fully developed pelvic girdles and lateral pelvic spines, partial or even complete pelvic reduction is common in freshwater populations (Blouw and Boyd 1992; Shapiro *et al*. 2004, 2006; Herczeg *et al*. 2010; Klepaker *et al*. 2013). Several factors may contribute to this, including the absence of gape-limited predatory fish and limited calcium availability, as well as the presence of certain insect predators (Reist *et al*. 1980; Reimchen *et al*. 1983; Giles 1983; Bell *et al*. 1993; Karhunen *et al*. 2013; Chan *et al*. 2010) in freshwater environments. Collectively, sticklebacks provide an important model system to study the genetic mechanisms underlying the adaptive parallel pelvic reduction at both intra- and inter-specific levels, under a wide range of population demographic settings. However, studies of parallel patterns of marine-freshwater divergence in nine-spined sticklebacks are still scarce (Herczeg *et al*. 2010; Shikano *et al*. 2010b; Wang *et al*. 2020), especially at the genetic level, precluding any comprehensive and conclusive comparison of the two species.

The genetic basis of pelvic reduction in the three-spined stickleback is well understood. Quantitative trait locus (QTL) mapping studies have identified a single chromosomal region containing the gene *Pituitary homeobox transcription factor 1* (*Pitx1*) that explains more than two thirds of the variance in pelvic size in crosses between individuals with complete pelvic girdles and spines, and pelvic-reduced individuals (Cresko *et al*. 2004; Shapiro *et al*. 2004; Coyle *et al*. 2007). Pelvic loss in the marine-freshwater three-spined stickleback model system (as well as in benthic-limnetic pairs of three-spined sticklebacks; Peichel *et al*. 2001) is predominantly caused by expression changes of the *Pitx1* gene due to recurrent deletions of the pelvic enhancer (*Pel*) upstream of *Pitx1* (Chan *et al*. 2010; Xie *et al*. 2019; in lake-stream pairs of three-spined sticklebacks, the genetic architecture of pelvic reduction is more diversified; Peichel and Marques 2017; Deagle *et al*. 2012; Stuart *et al*. 2017; Rennison *et al*. 2019). While pelvic reduction in freshwater is much less common in nine-than three-spined sticklebacks (Klepaker *et al*. 2013; Fig. 1), two independent QTL studies also identified *Pitx1* in linkage group (LG) 7 as a major cause for pelvic reduction in a Canadian (Shapiro *et al*. 2006) and a Finnish (Shikano *et al*. 2013) population. Another large effect region in LG4 was found to be associated with pelvic reduction in an Alaskan population (Shapiro *et al*. 2009). Similar to the three-spined stickleback, pelvic spine and pelvic girdle sizes are strongly correlated in the population from the Finnish pond Rytilampi, since *Pitx1* controls both phenotypes (Shikano *et al*. 2013; Fig. 1 and Supplementary Table 1). In contrast, a considerable amount of phenotypic variation with respect to these traits and their inter-correlations exists among different freshwater pond populations in northern Europe (Herczeg *et al*. 2010; Karhunen *et al*. 2013, 2014; Fig. 1). Given their high heritability (Blouw and Boyd 1992; Leinonen *et al*. 2011), the lack of correlation between spine and girdle lengths suggests that they can be independently controlled by different QTL. Thus, the genetic underpinnings of pelvic reduction in nine-spined sticklebacks (when it occurs) appear to be more diversified than those in the marine-freshwater three-spined stickleback model system (Merilä 2013, 2014).

**Figure 1.**
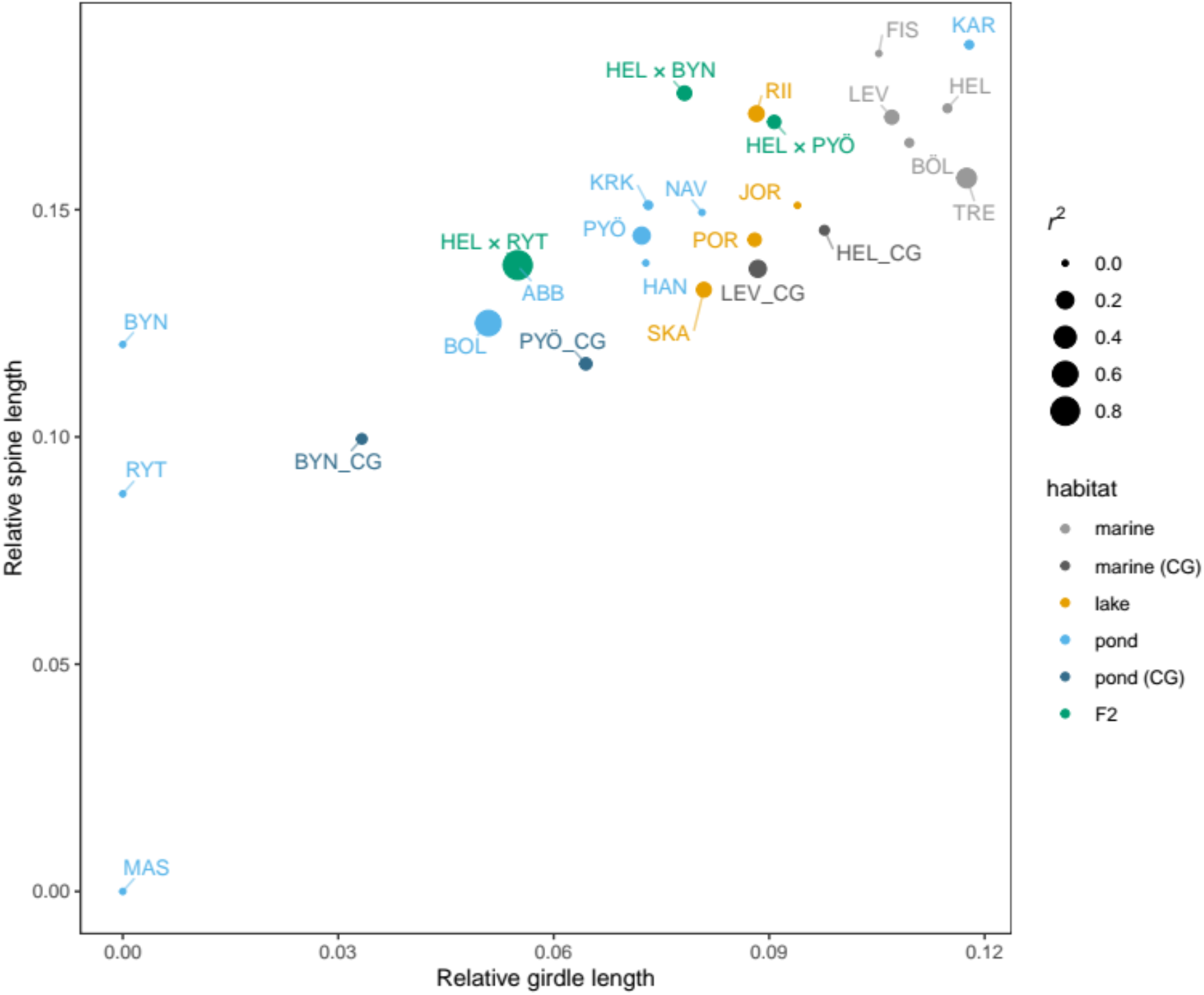
Summary of previously published nine-spined stickleback pelvic phenotypes from the wild and common garden experiments. Scatterplot depicts relative (to standard body length) pelvic spine and girdle lengths colored according to population and habitat type and with size indicating Pearson’s product moment correlation (*r^2^*) between spine and girdle lengths for a given population. Populations with complete spine and/or girdle reductions (BYN, RYT and MAS) are here given *r^2^*=0. Further details can be found in Supplementary Fig. 3 and Supplementary Table 1. Phenotypic data for F_2_ individuals from the current study are also presented (HEL × RYT, HEL × BYN, and HEL × PYÖ) for comparison. Details of sample locations and data collection can be found from Herczeg *et al*. (2010) and Karhunen *et al*. (2013). Data from common garden experiments is indicated by the suffix “_CG” in the x-axis labels.

Starting with a survey of previously published phenotypic data on pelvic reduction in the wild, we aimed to investigate the possible genetic heterogeneity underlying pelvic reduction in different nine-spined stickleback populations by mapping QTL for pelvic reduction in three independently colonised ponds. One was the previously studied Rytilampi (earlier analysed only with microsatellites, Shikano *et al*. 2013), now re-analysed along with two new populations (Bynästjärnen and Pyöreälampi) using >75 000 SNPs. These data were subjected to a cutting-edge mapping approach (Li Z. *et al*. 2017, 2018) that can provide more information on the source of the QTL effects than has been previously possible. To further assess the extent to which *Pel* could be responsible for pelvic reduction in the nine-spined stickleback, we screened the whole genomes of individuals from 27 wild populations for deletions in the genomic region spanning the *Pel* element and the *Pitx1* gene. Finally, utilising more comprehensive geographic sampling than in previous studies (Shikano *et al*. 2010a; DeFaveri *et al*. 2012; Merilä 2013), along with high-quality SNP data, we re-assessed the differences in population structuring and genetic isolation by distance (IBD) between nine-and three-spined sticklebacks. We hypothesised that, in contrast to three-spined sticklebacks, both the scarcity and variability in the genetic architecture of pelvic reduction in nine-spined sticklebacks is due to heterogeneity in the spatial patterns of standing genetic variation caused by strong population structuring – both among adjacent pond populations as well as in the ancestral sea population(s). Building on previous theoretical work, this hypothesis was tested using simulated data parameterised on population demographic parameters obtained from the empirical data.

## Materials and Methods

### Fish collection, crossing, and rearing

For the QTL crosses, three different marine F_0_ generation females from the Baltic Sea (Helsinki, Finland; 60°13’N, 25°11’E) were crossed with a freshwater F_0_ generation male from either Bynästjärnen (64°27’N, 19°26’E), Pyöreälampi (66°15’N, 29°26’E) or Rytilampi (66°23’N, 29°19’E) ponds. Fish crossing, rearing, and sampling followed the experimental protocol used in the earlier study of the Rytilampi population (Shikano *et al*. 2013; Laine *et al*. 2013). For Rytilampi, the F_0_ generation fish were artificially mated in the lab during the early breeding season of 2006 (Shikano *et al*. 2013), and the resulting full-sib F_1_-offspring were group-reared in aquaria until mature, as explained in Shikano *et al*. (2013). Two randomly chosen F_1_ individuals were mated repeatedly (seven different clutches) to produce the F_2_ generation between September and October 2008. The offspring were reared in separate aquaria. The same procedure was followed for Pyöreälampi (F_0_ mating: Jun 2011; F_1_ mating: Jul–Sep 2012; F_2_ rearing: Jul 2012–Apr 2013; 8 different clutches) and Bynästjärnen (F_0_ mating: Jun 2011; F_1_ mating: Nov 2013–Jan 2014, F_2_ rearing: Nov 2013–Aug 2014; 6 different clutches). The F_2_ fish were euthanized at 187, 238, and 238 days post-hatch for Rytilampi, Pyöreälampi and Bynästjärnen, respectively. At this point, the fish were on average 52.3 mm in standard length (Rytilampi = 48.6 mm; Pyöreälampi = 52.3 mm; Bynästjärnen = 53.6 mm), and weighed on average 1.34 g (wet weight; Rytilampi = 1.07 g; Pyöreälampi = 1.49 g; Bynästjärnen = 1.44 g). In total, 308, 283 and 279 F_2_ offspring were available for analyses from Helsinki × Bynästjärnen (HEL × BYN), Helsinki × Pyöreälampi (HEL × PYÖ) and Helsinki × Rytilampi (HEL × RYT) crosses, respectively.

The experiments were conducted under licenses from the Finnish National Animal Experiment Board (#STH379A and #STH223A). Experimental protocols were approved by the ethics committee of the University of Helsinki, and all experiments were performed in accordance with relevant guidelines and regulations.

### Morphological data

To visualise bony elements, all of the F_2_-progeny were stained with Alizarin Red S following Pritchard and Schluter (2001). Spine lengths was measured from its distal point to the basal point and the length of branch and girdle was measured at its maximum linear length with digital callipers to the nearest 0.01 mm (both sides of the body). Landmark definitions with illustrations depicting them can be found from Herczeg *et al*. (2010) and Shapiro et al. (2004). Although it is known that the left-right asymmetry of the pelvic girdle is heritable in sticklebacks (Blouw and Boyd 1992; Bell *et al*. 2007; Coyle *et al*. 2007), we did not specifically study this. To reduce the number of tests, the mean of the left and right-side measurements was used (analyses conducted for the two sides separately always yielded qualitatively similar results as the tests conducted with the averages; results not shown). All measurements were made by the same person twice with repeatability >0.97 for all traits (R; Becker 1984). The QTL analyses were performed on both absolute and relative (scaled to the total body length) trait values, but for all of the analyses that compared phenotypic data between populations (which also differ in body sizes), only relative trait values were used. Previously published phenotypic data from 19 wild populations were obtained from Herczeg *et al*. (2010) and Karhunen *et al*. (2013). These included data on pelvic spine and girdle lengths of wild-collected individuals from ten pond populations (Abbortjärn = ABB, Bolotjone = BOL, Karilampi = KAR, Kirkasvetinen lampi = KRK, Mashinnoje = MAS, Lil-Navartjärn = NAV, Hansmytjärn = HAN, Rytilampi = RYT, Bynästjärnen = BYN), four lake populations (Iso Porontima = POR, Riikojärvi = RII, Joortilojärvi = JOR, Västre-Skavträsket = SKA) and five marine populations (Fiskebäckskil = FIS, Trelleborg = TRE, Bolesviken = BÖL, Helsinki = HEL LEV = Levin Navolak), as well data on common garden-reared F_1_ generation individuals from two marine (HEL and LEV) and two pond (BYN and PYÖ) populations. Visibly broken spines were treated as missing data. A map showing the geographic location of these populations is provided in Supplementary Figure 1.

### Genotyping and linkage map construction

The RAD sequencing protocol used to obtain the SNP data was the same as in Yang *et al*. (2016) and Li Z. *et al*. (2017). In short, genomic DNA was extracted from ethanol preserved fin clips using the phenol-chloroform method (Taggart *et al*. 1992). DNA was fragmented with PstI restriction enzyme and the resulting 300–500 bp long fragments were gel purified. Illumina sequencing adaptors and library specific barcodes were ligated to the digested DNA fragments, and the barcoded RAD samples were pooled and sequenced on 24 lanes of the Illumina HiSeq2000 platform with 45 bp single-end strategy. RAD library construction and sequencing were conducted by BGI HONGKONG CO., LIMITED. After eliminating adapters and barcodes from reads, a quality check was done using FastQC (http://www.bioinformatics.bbsrc.ac.uk/projects/fastqc/).

A linkage map based on the three crosses was constructed using Lep-MAP3 (LM3; Rastas 2017), as described in detail in Li H. *et al*. (2018). LM3 can infer the parental/grandparental phase based on dense SNP data, which allowed us to utilise the four-way cross QTL mapping method described below. Input data were generated by first mapping individual fastq files to the nine-spined stickleback reference genome using BWA mem (Li H. 2013) and SAMtools (mpileup; Li H. *et al*. 2009), followed by pileupParser.awk and pileup2posterior.awk scripts from the LM3 pipeline using default parameters (see Supplementary File 1 for details).

### Dimensionality reduction by linkage disequilibrium network clustering

It is essential in QTL mapping to correct for multiple testing in order to reduce the rate of false positives. Moreover, in large genomic datasets, physically adjacent SNPs – particularly those from experimental crosses – are often in linkage disequilibrium (LD), i.e. correlated. Since a group of SNPs in high LD explains similar amounts of genetic variation in a given trait, it is reasonable to apply a dimensional reduction procedure before QTL-mapping to exclude the redundant information from the data. Here we used linkage disequilibrium (LD) network clustering (LDn-clustering) and PC regression as dimensionality reduction tools prior to single-locus QTL mapping (Li Z. *et al*. 2018). The first step of this approach involves an extension of a method developed by Kemppainen *et al*. (2015), which uses LD network analysis for grouping loci connected by high LD. The second step involves principal component analysis (PCA) as a method for dimensionality reduction in each cluster of loci connected by high LD (LD-clusters). This was achieved by the function LDnClustering in the R-package LDna (v.0.64; Li Z. *et al*. 2018). The method used here differs slightly from the original method described in Li Z. *et al*. (2018) to increase efficiency in both computational speed and complexity reduction (see Supplementary File 2 for details).

### QTL mapping: four-way crosses model

In some circumstances, such as a four-way cross (Xu 1996), F_1_ hybrids of two heterozygous parents (Van Ooijen 2009), and an outbred F_2_ design (Xu 2013a), it is possible that up to four different alleles – A and B from the dam, and C and D from the sire – segregate in the population. In such cases, a linear regression model for QTL analysis of the outbred F_2_ data (Li Z. *et al*. 2018) is defined by:

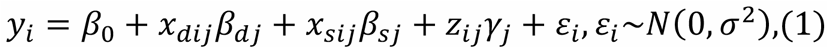

where *y_i_* is the phenotype value of individual *i* (*i* = 1,…, *n*), *x_dij_*, x_*sij*_, *z_ij_* are genotypes coded as

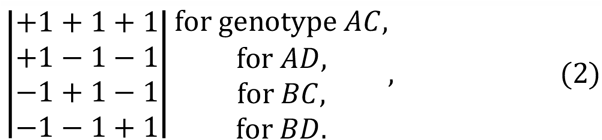

(Xu 2013b), *β_0_* is the parameter of the population mean, *β_dj_* is the substitution effect of alleles A and B of the dam (i.e. the grandfather in F_0_) at the locus *j* (*j* = 1,…, *p*; *p* is the total number of SNPs), *β_dj_* is the substitution effect of alleles C and D of the sire (i.e. the grandmother in F_0_), *γ_j_* is the dominance effect, and *ε_i_* is the residual error term mutually following an independent normal distribution.

The model (1) requires the knowledge of the grandparental phase (produced by LM3) with the benefit that the source (*viz*. F_0_ female or F_0_ male) of the observed QTL effect can be inferred, as described in more detail in Supplementary File 3. A multiple correction on the basis of permutation tests was further conducted to control for false positives due to multiplicity (Li *Z*. *et al*. 2017) with 1×10^5^ replicates.

### Estimating the proportion of phenotypic variance explained by SNPs

The overall proportion of phenotypic variance (PVE) explained jointly by all SNPs (an approximation of the narrow sense heritability) was obtained using LASSO regression (Tibshirani 1996), which incorporates all the SNPs into a multi-locus model:

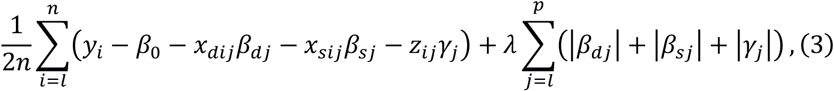

where the *l*_1_ penalty term 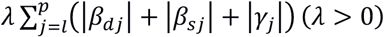 shrinks the regression parameters towards zero; all other symBÖLs are defined in the same way as in Equation (1).

Following Sillanpää (2011), the PVE can be estimated by the formula:

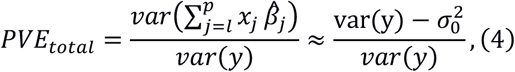

where 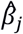 is the effects of the SNPs, and 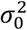 is the LASSO residual variance estimated by a cross-validation-based estimator introduced by Reid *et al*. (2016). The PVE explained by each linkage group was estimated on the basis of the LASSO estimates using the following formula:

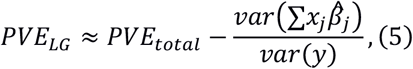

where 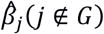 represents all the effects estimated by the LASSO of the SNPs that do not belong to a set of SNPs (e.g. to a chromosome/linkage group). A similar approach was used to estimate the contribution of grandparental alleles and to evaluate the dominance component by treating the coding systems [*x_dij_, x_sij_, z_ij_*] (2) as different groups of SNPs.

### Scanning for Pel deletions in full-genome sequence data

A minimum of 20 samples from populations RYT, MAS, BÖL, BYN and PYÖ, and 10–31 samples from an additional 22 populations (from Northern Europe and USA; Supplementary Fig. 1 and Supplementary Table 2) were sequenced to 10× coverage by BGI HONGKONG CO., LIMITED. Reads were mapped to the nine-spined stickleback reference genome (Varadharajan *et al*. 2019) with BWA mem (Li H. 2013), and site-wise sequencing coverages were computed with SAMtools (depth; Li H. *et al*. 2009). Relative sequence depths across the *H2afy-Pitx1* intergenic region were estimated for the five focal populations by first computing the median depths for 1000 bp sliding windows, and then normalising these by the median depth for the full intergenic region. The Pel-2.5kb^SALR^ region was extracted from the original BAC contig (GenBank accession number GU130433.1) and mapped to the nine-spined stickleback intergenic region with minimap2 (Li H. 2018). The sequencing depths for the Pel-2.5kb^SALR^ were normalised by dividing the mean depths of the *Pel* region by the mean depths of the full intergenic region. Gene annotations and relative sequencing depths (average and confidence intervals) were computed and visualised using the R-package Gviz (Hahne and Ivanek 2016). Lastly, we scanned the literature for genes that are known to affect pelvic and hind limb development, and searched for potential matches in the nine-spined stickleback genome (Varadharajan *et al*. 2019) in regions where significant QTL were found.

### Simulations

The repeated local adaptation in independently colonised freshwater stickleback populations is widely considered to be due to selection on standing genetic variation available in the sea, which in turn is maintained by recurrent gene flow from previously colonised freshwater populations (the “transporter hypothesis”; Schluter and Conte 2009). The effects of standing genetic variation (assuming a panmictic ancestral population) on the probability of parallel local adaptation have already been studied (Ralph and Coop 2015a; MacPhearson and Nuisimer 2016; Lee and Coop 2017; Galloway and Cresko 2019), along with the effects of population structure on adaptation in continuous landscapes (Hermisson and Pennings 2005; Ralph and Coop 2010, 2015b). However, no simulations to date have explicitly considered the effects of heterogeneous patterns in the distribution of standing genetic variation on local adaptation/parallel evolution. Thus, we used forward-in-time simulations to explore the effects of the differences in population demographic parameters between nine- and three-spined sticklebacks on the probability of local adaptation, as described in Supplementary File 4. These simulations were parameterised using empirical estimates of population structure and isolation by distance in the sea for the two species (Supplementary File 5).

In short, two freshwater populations were connected to different marine populations at the ends of a stepping-stone chain of marine populations with the carrying capacity (K) and migration (M) both in the sea, and between the founding (marine) and founder (freshwater) population (Supplementary Fig. 2), thus generating comparable patterns of population structuring and IBD in the sea as in the empirical data (see Results). The two freshwater populations were colonised once and allowed to become locally adapted for 10,000 years (20,000 generations). All freshwater adaptation was due to standing genetic variation in the sea that was continually replenished from a “refuge” freshwater population (situated at an equal distance from the two focal freshwater populations), as well as from the two focal freshwater populations once they were colonised. Three different genetic architectures were studied. One (architecture A) corresponds to a single large effect QTL (allowing 100% local adaptation, but with minor effect QTL also affecting the trait), another (architecture B) to a situation where at least two loci are needed to allow local adaptation (allowing 40% or 60% local adaptation), and the third (architecture C) corresponds to the same scenario as A, except that the large effect locus is recessive. Mutation rate was fixed at 1e^-8^ and all loci were polymorphic at the beginning of the simulations. More details are given in Supplementary File 4.

## Results

### Heterogeneity of pelvic reduction in the wild

Re-analysis of previously published phenotypic data from the wild confirmed a high degree of variation among different populations with respect to pelvic spine and girdle lengths and their inter-correlations (Fig. 1, Supplementary Table 1 and Supplementary Fig. 3). For instance, while spines were absent and pelvic girdles strongly reduced (but not completely absent) in RYT, spines and girdles were completely lacking in the MAS population (Fig. 1 and Supplementary Table 1). Furthermore, in the BYN population, spines were absent but pelvic girdles were only partially reduced; in BÖL (a population adjacent to MAS), large variation in both spine (SD = 0.037) and girdle (SD = 0.047) lengths was observed, although these two traits were strongly correlated (*r^2^* = 0.61, Fig. 1, Supplementary Table 1 and Supplementary Fig. 3). This suggests that a large effect locus affecting both pelvic spines and girdles segregated in this population at the time of sampling. In six pond populations (*viz*. ABB, KAR, KRK, NAV, HAN, and PYÖ; Fig. 1), relative spine (mean = 0.079) and girdle (mean = 0.15) lengths were only slightly smaller relative to the marine populations (0.11 and 0.16 for spine and girdle lengths, respectively; Supplementary Table 1). A lack of pelvic reduction (relative to the marine populations) was observed only in one pond population (KAR; Fig. 1, Supplementary Table 1 and Supplementary Fig. 3).

### QTL mapping of pelvic reduction in the Helsinki × Rytilampi cross

After LD-network based complexity reduction, all QTL mapping analyses were performed on 241 PCs (six sex-linked PCs were removed). Re-analyses of the 279 F_2_ progeny from the HEL × RYT cross confirmed a single QTL region on LG7 for both pelvic spine and girdle lengths in the single-locus analyses (Fig. 2a, b and Table 1). In the multi-locus analyses (absolute trait values), LG7 explained 74–86% of the PVE for both spine and girdle lengths, while all other chromosomes individually explained less than 3% of the phenotypic variance (Table 2). An approximately equal amount of the phenotypic variance was explained by alleles inherited from the F_0_ male (pond individual) and the F_0_ female (marine individual; ~30% of the total variance for all traits; Table 2), respectively, with 15–21% of the phenotypic variance also explained by dominance effects. This is the expected outcome for a recessive QTL when the F_0_ individuals are fixed for different large effect causal alleles, and when no additional smaller effect loci affect the trait (Klug and Cummings 2018). This is also consistent with a strong correlation between relative pelvic spine and girdle lengths (*r^2^* = 0.85, Fig. 1, Supplementary Fig. 3 and Supplementary Table 1).

**Figure 2.**
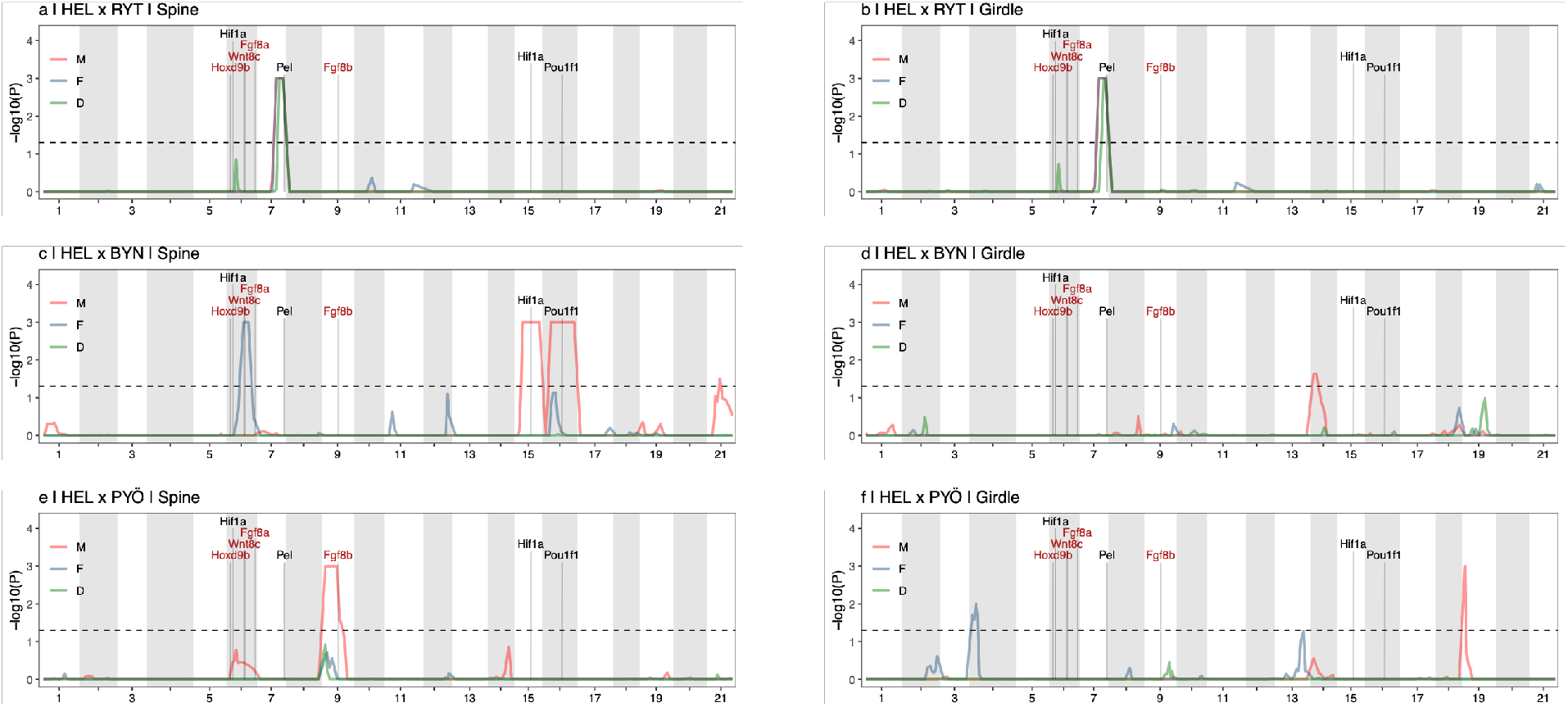
Quantitative trait locus mapping of pelvic reduction in three independent stickleback crosses. Single-mapping four-way analyses of four morphological traits associated with pelvic reduction in (a-b) HEL × RYT cross, (c-d) HEL × BYN cross, and (e-f) HEL × PYÖ cross. QTL for pelvic spine length and girdle length are shown, with the x-axis indicating position in centi Morgans (cM). Results are based on permutations, and the dotted vertical line indicates genome wide significance at α = 0.05. Results are shown separately for alleles inherited from the male F_1_ (M), the female F_1_ (F), together with the dominance effect (D) according to the legend. Absence of a dominance effect indicates that the trait inheritance is additive, whereas a peak only for M or F indicates that the allelic effect was segregating in the F_0_ male or the F_0_ female, respectively (see Supplementary File 3 for details). Candidate genes involved in pelvic development are indicated with black text representing genes that affect expression of *Pitx1*, and red text indicates genes that affect pelvic development downstream of *Pitx1* expression. Results for analyses based on relative trait values can be found in Supplementary Figure 5.

**Table 1.** Summary of QTL-mapping analyses. Each row corresponds to a significant (arbitrarily numbered) QTL region, with the proportion variance explained (PVE), jointly estimated for all PCs (one for each significant LD-cluster) extracted from such regions. Coding indicates whether the QTL was significant for the alleles inherited from the F_1_ female (♀), the F_1_ male (♂) or the dominance effect (dom). Effect size (*β*) is based on the first PC extracted from all SNPs from each significant QTL region. “Std.” indicates whether the trait was standardised (Yes) or not (No). *P* and *P_COR_* represent nominal and corrected *P*-values from single-mapping four-way analyses, respectively. Results for standardised trait values are only shown for QTL that were not also significant for absolute trait values.

| Cross | Trait | Coding | LG | QTL | Std. | $P$ | $P_{COR}$ | PVE<sub>TOT</sub> | $\beta$ |
| --- | --- | --- | --- | --- | --- | --- | --- | --- | --- |
| HEL × RYT | Spine | male | 7 | 1 | No | 1.31e-04 | 0.019 | 0.285 | 1.654 |
| HEL × RYT | Spine | female | 7 | 1 | No | 1.26e-04 | 0.026 | 0.311 | 1.573 |
| HEL × RYT | Girdle | male | 7 | 1 | No | 3.66e-04 | 0.04 | 0.311 | 2.041 |
| HEL × RYT | Girdle | female | 7 | 1 | No | 1.08e-04 | 0.024 | 0.263 | 1.711 |
| HEL × RYT | Spine | dom | 7 | 1 | No | 1.08e-07 | <0.001 | 0.213 | 1.432 |
| HEL × RYT | Girdle | dom | 7 | 1 | No | 1.35e-04 | 0.028 | 0.154 | 1.463 |
| HEL × BYN | Spine | male | 15 | 2 | No | 6.12e-06 | 0.001 | 0.095 | 0.54 |
| HEL × BYN | Spine | male | 16 | 3 | No | 1.16e-04 | 0.01 | 0.086 | 0.493 |
| HEL × BYN | Spine | male | 21 | 4 | No | 3.36e-04 | 0.031 | 0.037 | 0.36 |
| HEL × BYN | Spine | female | 6 | 5 | No | 1.37e-04 | 0.018 | 0.07 | 0.5 |
| HEL × BYN | Girdle | male | 14 | 6 | No | 2.49e-04 | 0.024 | 0.039 | 0.307 |
| HEL × PYÖ | Spine | male | 9 | 7 | No | 8.96e-11 | <0.001 | 0.108 | 0.352 |
| HEL × PYÖ | Girdle | male | 19 | 8 | No | 1.56e-05 | 0.001 | 0.056 | 0.325 |
| HEL × PYÖ | Girdle | female | 4 | 9 | No | 1.32e-04 | 0.02 | 0.045 | 0.301 |
| HEL × BYN | Spine | female | 16 | 10 | Yes | 2.53e-04 | 0.03 | 0.047 | 0.007 |
| HEL × BYN | Girdle | female | 1 | 11 | Yes | 2.13e-05 | 0.002 | 0.077 | 0.006 |
| HEL × BYN | Girdle | male | 1 | 11 | Yes | 1.41e-04 | 0.016 | 0.026 | 0.005 |
| HEL × PYÖ | Spine | male | 1 | 13 | Yes | 4.75e-06 | <0.001 | 0.075 | 0.005 |
| HEL × PYÖ | Girdle | male | 1 | 13 | Yes | 9.69e-05 | 0.016 | 0.044 | 0.005 |

### QTL mapping of pelvic reduction in the Helsinki × Bynästjärnen cross

Among the 308 F_2_ progeny of the HEL × BYN cross, single-locus mapping analyses of pelvic spine lengths detected two significant QTL on LG15 (PVE = 8.9%) and LG16 (PVE = 13.7%; Fig. 2c, Table 1 and 2) for alleles deriving from the F_0_ male (pond individual). Thus, the causal alleles for these QTL segregated in the F_0_ pond male (as explained in Supplementary File 3). No dominance effects were detected for these QTL, suggesting that the allelic effects were additive. One QTL on LG6 with an allelic effect deriving only from the F_0_ female was also detected (Fig. 2c and Table 1). The QTL significance profiles, in particular for LG15 and LG16, spanned large genomic regions with no obvious peaks (in contrast to LG7 in the HEL × RYT cross; Fig. 2a, b). This remained true when analysing all SNPs individually (fine-mapping; Supplementary Fig. 4). In addition, phenotypes of the F_2_ individuals from the HEL × RYT cross were not normally distributed and the individual and multi-locus phenotypic effects on spine lengths for the QTL on LGs 6, 15 and 16 indicated epistatic interactions (Supplementary File 6). One significant male QTL on LG4 for girdle length was also detected (Fig. 2d and Table 1). Results for relative trait values were highly similar to the absolute trait values, except for an additional significant male and female QTL on LG1, as well as another significant male QTL on LG14 (Supplementary Fig. 5, Table 1 and Supplementary Table 3).

**Table 2.** Proportion phenotypic variance explained (PVE) in pelvic traits. Percentages of total phenotypic variance explained by different linkage groups (LG), by all SNPs (Tot), by loci inherited from females (♀) and males (♂), as well as the dominance effect (Dom). Results are shown for each cross and trait separately, and for absolute trait values.

| LG | HEL x RYT |  | HEL x BYN |  | HEL x PYÖ |  |
| --- | --- | --- | --- | --- | --- | --- |
|  | Spine | Girdle | Spine | Girdle | Spine | Girdle |
| 1 | - | 0.01 | 0.03 | 0.05 | - | - |
| 2 | - | - | 0.01 | 0.03 | - | 0.02 |
| 3 | - | - | - | 0.01 | - | - |
| 4 | - | 0.02 | 0.01 | 0.01 | - | 0.03 |
| 5 | - | - | - | - | - | - |
| 6 | - | - | 0.03 | - | 0.02 | - |
| 7 | 0.85 | 0.82 | 0.02 | 0.02 | - | - |
| 8 | - | - | 0.01 | 0.01 | - | - |
| 9 | - | - | - | 0.02 | 0.11 | - |
| 10 | - | - | - | 0.06 | - | - |
| 11 | - | - | 0.02 | - | - | - |
| 12 | 0.02 | 0.02 | 0.03 | 0.01 | 0.01 | - |
| 13 | - | - | - | 0.01 | - | 0.02 |
| 14 | - | - | - | 0.05 | 0.01 | 0.02 |
| 15 | - | - | 0.09 | 0.01 | - | - |
| 16 | - | - | 0.12 | 0.01 | - | - |
| 17 | - | - | 0.01 | 0.01 | - | - |
| 18 | - | - | - | 0.04 | - | 0.01 |
| 19 | - | - | 0.02 | 0.06 | - | 0.05 |
| 20 | - | - | - | 0.01 | - | - |
| 21 | - | - | 0.03 | - | 0.01 | - |
| **Tot** | 0.8 | 0.76 | 0.39 | 0.32 | 0.14 | 0.16 |
| ♂ | 0.3 | 0.38 | 0.24 | 0.16 | 0.14 | 0.08 |
| ♀ | 0.29 | 0.27 | 0.17 | 0.06 | - | 0.09 |
| **Dom** | 0.23 | 0.17 | 0.01 | 0.13 | 0.01 | - |

Multi-locus analyses (for absolute trait values) identified 11 LGs that accounted for at least 3% of the phenotypic variance in pelvic spine or girdle lengths in the HEL × BYN cross (Table 2). The largest of these effects were found on LG15 and LG16, which explained 9% and 12% of the variation in pelvic spine lengths, respectively (Table 2). For pelvic spine and girdle lengths, 39% and 32% of the PVE, respectively, were accounted for by all SNPs in the dataset, which equates to the narrow sense heritability (*h^2^*) also accounting for dominance (but not epistatic interaction) effects. For spine lengths, 24% of the total PVE was attributed to alleles deriving from the F_0_ male, and 17% was attributed to alleles deriving from the F_0_ female; only 1% was attributed to the dominance effect (Table 2). For girdle lengths, the corresponding numbers were 16%, 6% and 13% (Table 2).

Compared to the HEL × RYT cross, the correlation between relative pelvic spine and girdle lengths was much weaker in HEL × BYN cross (*r^2^* = 0.11, Fig. 1, Supplementary Fig. 3 and Supplementary Table 1) consistent with pelvic spine and girdle reductions being independently controlled by different QTL. Furthermore, of the 306 F_2_ individuals, only four displayed complete lack of spines (and the spine was absent in the F_0_ male), consistent with spine length being controlled by multiple additive loci.

### QTL mapping of pelvic reduction in the Helsinki × Pyöreälampi cross

In the HEL × PYÖ cross (283 F_2_ individuals), one QTL for spine length was found on LG9, which was explained by alleles inherited from the F_0_ male. Two significant QTL for girdle length were found, one on LG19 (explained by alleles inherited from the F_0_ male) and the other on LG4 (explained by alleles inherited from the F_0_ female; Fig. 2f and Table 1). In the multi-locus analyses, the PVE for different LGs mirrored these results; the LGs that contain significant QTL explain most of the PVE, while PVE for all other LGs were <4% (Table 2). Overall, the multi-locus analyses revealed that PVE for pelvic traits was lower than in the HEL × BYN cross; 14% and 16% for spine length and girdle length, respectively, with <2% PVE due to dominance effects (Table 2). When analysing relative trait values, one additional female QTL peak was found for both girdle and spine lengths on LG1 (Supplementary Fig. 5, Table 1 and Supplementary Table 3). Due to the large size of the identified QTL regions, it is not possible to know whether this QTL and that on LG1 detected in the HEL × BYN cross are due to the same or different underlying causal mutations (we consider this as a single QTL region).

### Scanning for Pel deletions in the full-genome sequence data

In the full-genome re-sequencing data of wild-collected individuals, a large deletion upstream of *Pitx1* was fixed for all 21 individuals from Rytilampi, where pelvic reduction maps to this region (Fig. 3). The deletion is around 3.5 kb (LG7:17015109-17018744; Supplementary Fig. 6) in size and fully encompasses the Pel-2.5kb^SALR^ region of the three-spined stickleback (Chan *et al*. 2010). No comparable deletion was found in any other individuals in the dataset (Fig. 3b). This included the two White Sea populations, in which either complete reduction of both the pelvic spines and girdles was observed (MAS), or a putative large effect locus affecting both spine and girdle length was found to be segregating (BÖL; Fig. 1; Supplementary Fig. 3 and Supplementary Table 1).

**Figure 3.**
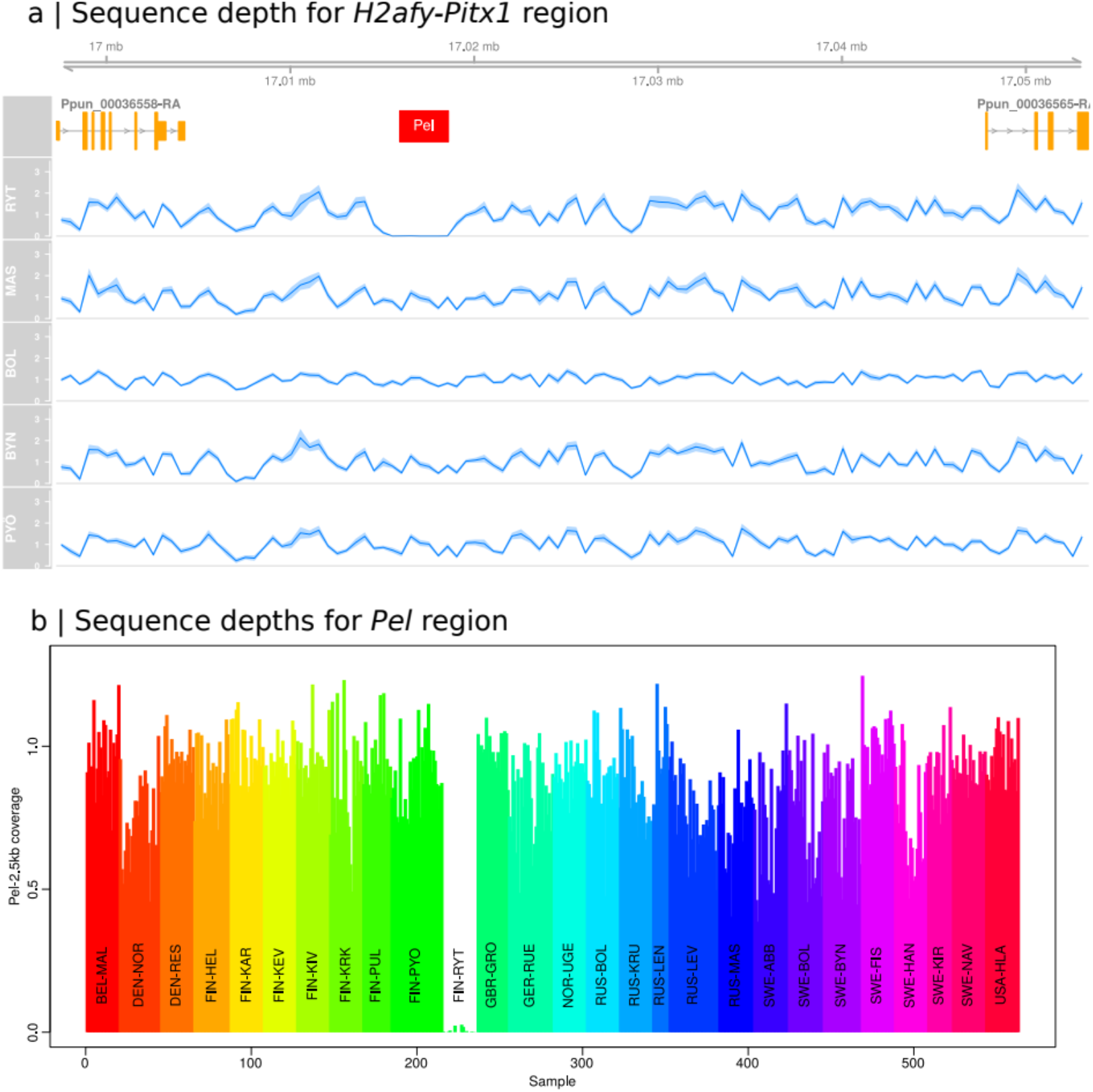
Sequence depths for *H2afy-Pitx1* region. (a) Relative sequencing depth across the *H2afy-Pitx1* intergenic region for 10 individuals from five different populations. Blue line and shading indicate population average and 95% confidence intervals, respectively. The 3.5kb deletion in RYT, seen as a dip in sequencing depth, fully encloses the Pel-2.5kb^SALR^ of three-spined stickleback indicated by the red box. (b) Normalised sequencing depths for the Pel-2.5kb^SALR^ region for 563 samples from 27 populations, showing that the complete deletion of the *Pel* region is unique to RYT. Phenotypic data in the Russian populations MAS and BÖL suggest that a large effect locus is responsible for pelvic reductions, and BYN and PYÖ (analysed here) are populations in which pelvic reduction does not map to LG7.

### Candidate genes

Seven candidate genes or regulatory elements for pelvic reduction identified from the literature (Supplementary Table 4) were found in the LGs with significant QTL (Fig. 2). *Hif1a* (Mudie *et al*. 2014), *Pel* (Chan *et al*. 2010) and *Pou1f1* (Kelberman *et al*. 2009) are known to regulate the expression of *Pitx1*, whereas four genes (*Fgf8, Wnt8c, Wnt8b* and *Hoxd9*) are involved in the pelvic fin/hind limb development downstream of *Pitx1* (Don *et al*. 2012; Tanaka *et al*. 2005). However, aside from *Pel* (LG7), only three – *Wnt8c* (LG6), *Hif1a* (LG15) and *Pouf1f1* (LG16) – were clearly within the significant QTL regions (Fig. 2). One candidate locus, *Hif1a*, is also on LG1 where significant QTL peaks were found when analysing relative (but not absolute) trait values (Table 1 and Supplementary Fig. 5).

### Simulations

In the empirical data from the marine populations, IBD (the slope of the regression line between linearised *F_ST_* and geographic distance point distance in km) was 2.13^-7^ (95% of the 1e^6^ bootstrap replicates were between 2.05^-7^ and 2.2^-7^) for nine-spined sticklebacks, and 1.53^-8^ (95% of the bootstrap replicates were between 1.10^-8^ and 1.96^-8^) for three-spined sticklebacks. Thus, the slope of the IBD regression line in this dataset was 13.9 times higher for nine-spined sticklebacks compared to three-spined sticklebacks (Fig. 4a). The geographic distance (which is on an arbitrary scale) in the simulated data was scaled based on the observed levels of IBD in the empirical data. This resulted in the distance between the two simulated marine populations furthest away from each other (i.e. the marine populations from which the focal freshwater populations were founded) corresponding to 264 km and 352 km for three- and nine-spined sticklebacks, respectively (Fig. 4a). Thus, with comparable levels of IBD as in the empirical data, our simulations mimic the levels of parallel evolution that can be expected in three- and nine-spined sticklebacks at relatively short geographic scales (<400 km). The difference in IBD between the two species is also close to that in the empirical data (264/352 km = 0.75). In the empirical data, genetic differentiation between freshwater habitats for populations <400 km from each other (mean *F_ST_* = 0.19 and 0.49 for three- and nine-spined sticklebacks, respectively) was also on par with the simulations (mean [across simulation replicates] *F_ST_* = 0.21 and 0.58, respectively). Notably, *F_ST_* was >0.8 (linearised *F_ST_*>4) between Pyöreälampi (no pelvic reduction) and Rytilampi (pelvic reduction controlled by *Pitx1/Pel*), although these ponds are situated only 15 km from each other (Fig. 4a; max and median *F_ST_* = 0.96 and 0.62, respectively, for all pairwise freshwater-freshwater comparisons). This is higher than the *F_ST_* between any pair of three-spined stickleback populations in the data (max = 0.78 and median = 0.26 for all pairwise freshwater-freshwater comparisons).

**Figure 4.**
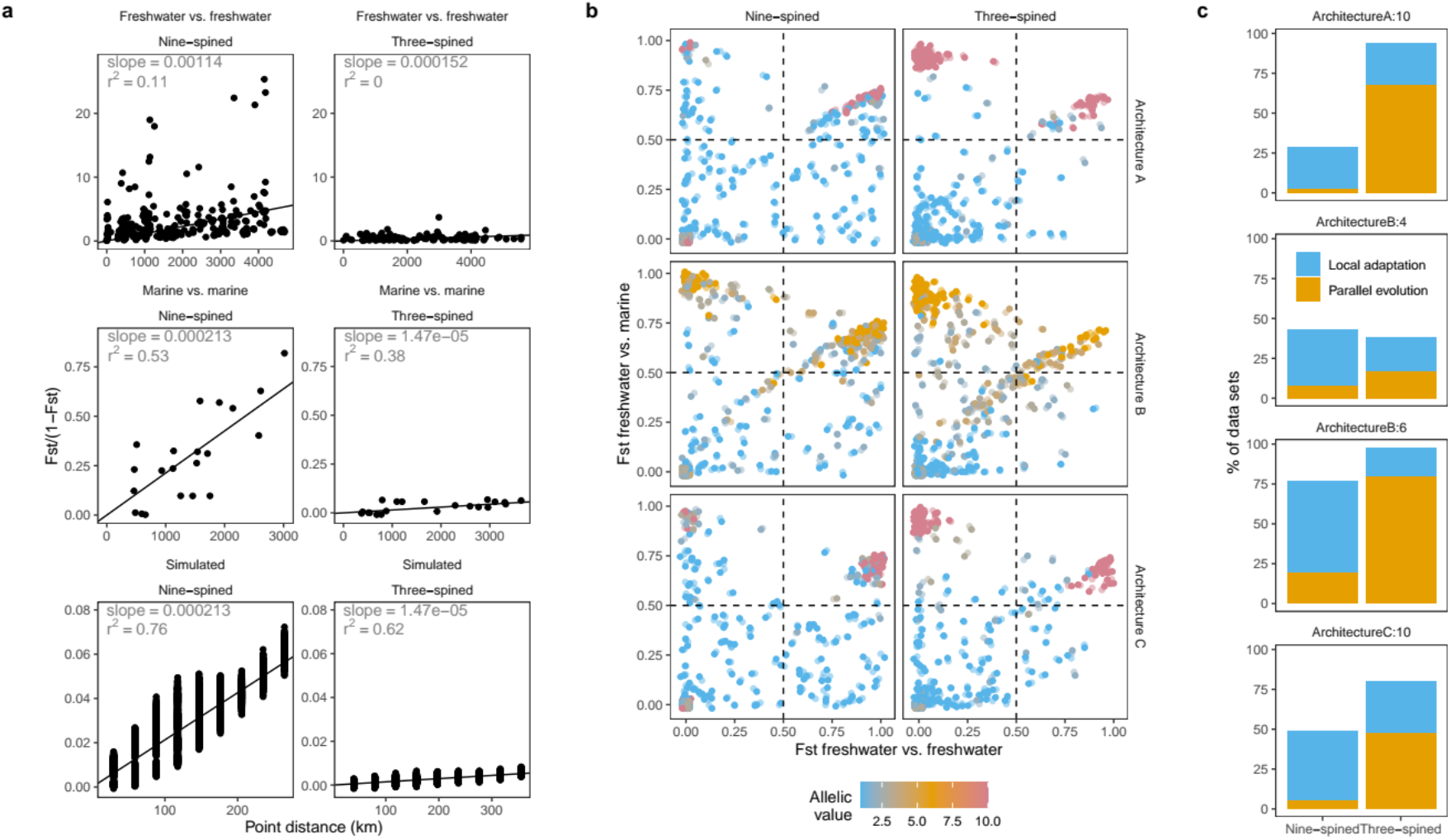
Simulation results. (a) depicts linearised *F_ST_* against geographic distance (IBD) for empirical and simulated data, with slope and squared Pearson’s product moment correlation coefficient indicated. Geographic distance for simulated data (IBD in the sea) is scaled to match the slope for the IBD-plot in the sea in the empirical data. (b) depicts freshwater-freshwater *F_ST_* against marine-freshwater *F_ST_* from all QTL from all simulated data (n = 100), with allelic effect sizes indicated as shown in legend (phenotypic optima in marine and freshwater habitats are 0 and 20, respectively). Loci in the upper left quadrant are classified as being involved in parallel evolution, and loci in the upper right quadrant are loci that are involved in local adaptation in only one freshwater population. This data is summarised in (c) focusing on the four largest effect loci, with genetic architecture and effect sizes indicated by the figure titles.

### Do population demographic parameters influence local adaptation?

In the simulations (Supplementary File 4), the relationship between marine-freshwater *F_ST_* and freshwater-freshwater *F_ST_* depended on both the species and genetic architecture (Fig. 4b). For instance, when the genetic architecture included one additive large effect locus (architecture A; homozygotes for this locus were 100% locally adapted to freshwater), this locus was often involved in parallel evolution when population structure was parameterised based on three-spined sticklebacks (65% of replicates, n = 100), but not when population structure was parameterised based on nine-spined sticklebacks (3% of the replicates). In 20% of the replicates for both species, the freshwater allele for this locus was fixed in only one of the focal freshwater populations (i.e. local adaptation, but not parallel evolution). When the trait under selection was controlled by several medium effect loci (architecture B), parallel evolution was more common in both stickleback scenarios, particularly for the locus with the largest allelic value (20% and 80% for nine- and three-spined sticklebacks, respectively; Fig. 4b, c; homozygotes for this locus were 60% locally adapted to freshwater). Local adaptation also occurred in 57% of the replicates in the nine-spine stickleback-like scenario, and in 18% in the 3-sp (Fig. 4b, c). For the locus with the second highest allelic value in architecture B (homozygotes for this locus were 40% locally adapted to freshwater), the cases of both parallel evolution and local adaptation collectively dropped to 38% and 43% three and nine-spined like stickleback scenarios, respectively. With one non-additive large effect locus with recessive alleles locally adapted to freshwater (architecture C), this locus was less likely to be involved in parallel evolution in the three-spined stickleback-like scenario (48%), compared to architecture A. However, results for the nine-spined stickleback-like scenario was similar to architecture A (6% parallel evolution; Fig. 4b, c). Thus, at relatively short (<400 km) geographical distances, parallel evolution is an expected outcome in three-but not nine-spined stickleback-like scenarios, particularly when a single additive large effect locus is responsible for freshwater adaptation. Results addressing the questions of how local adaptation depend on ancestral allele frequency are presented in Supplementary File 7 and Supplementary Figure 7, and QTL-use in parallel evolution as a function of allelic effect size is shown in Supplementary Figure 8.

## Discussion

The results demonstrate that pelvic reduction in nine-spined sticklebacks is not nearly as common as in three-spined sticklebacks, and when it does exist, the genetic basis is more variable. This is consistent with the re-analyses of neutral population genomic data showing that both IBD in the sea and the genetic structure among marine populations are >10 times higher in nine-spined sticklebacks, causing heterogeneity in the distribution of standing genetic variation that is not present to the same extent in the marine three-spined stickleback populations. Thus, the level and distribution of ancestral variation available for local adaptation (and also for parallel phenotypic evolution) is likely a function of population demographic parameters, with local adaptation being less likely in poorly connected species, as suggested by Merilä (2013, 2014) and corroborated by our simulations. However, other non-mutually exclusive factors, such as the genetic architecture (i.e. dominance effects, heritability, mutation rates and numbers of causal loci involved), as well as non-parallelism in phenotypic selection optima are also likely to play roles. In the following, we discuss the possible causes of the discrepancy in pelvic structure development between nine- and three-spined sticklebacks, as well as their implications to our understanding of adaptive evolution in the wild.

### Can the genetic architecture of pelvic reduction be explained by population demographic parameters?

Together with earlier QTL studies (Shapiro *et al*. 2006, 2009; Shikano *et al*. 2013), we show that multiple genomic regions (11 QTL, ten of which are novel to this study) are associated with pelvic reduction in nine-spined sticklebacks across their distribution range. Only one small effect QTL region (LG1; Table 1 and Supplementary Fig. 5) was shared between any two crosses (HEL × BYN and HEL × PYÖ) in our study, but even here it is not certain whether the underlying causal mutations are the same. Although the high frequency of *Pel* deletions (disrupting pelvic armour development) means that most of the deletions associated with pelvic reduction in three-spined sticklebacks are independently derived (Xie *et al*. 2019), *Pitx1/Pel* is nevertheless predominantly responsible for pelvic reduction in three-spined sticklebacks (Chan *et al*. 2010; Xie *et al*. 2019). In contrast, the major effect and recessive *Pitx1/Pel* allele in nine-spined sticklebacks is known to be responsible for pelvic reduction in only one Canadian population and Rytilampi. In both Bynästjärnen and Pyöreälampi, pelvic reduction is less heritable, polygenic and additive. In addition, based on morphological data, it seems likely that major effect loci not associated with *Pel*-deletions (Fig. 3) control pelvic reduction in two Russian ponds, with one additional Alaskan population being controlled by a large effect additive locus mapping to LG4 (Shapiro *et al*. 2009). We do not yet know the mutation rate of the *Pel*-deletions in nine-spined sticklebacks so naturally, the lack of *Pel*-deletions in nine-spined sticklebacks could be simply due to lower mutation rates compared to three-spined sticklebacks. Regardless, pelvic reduction in the nine-spined stickleback is much less common than in three-spined sticklebacks in general (Klepaker *et al*. 2013), and when it occurs, the QTL that control pelvic reduction are much more variable with respect to heritability, effect size distribution and dominance relationships. Assuming that selection for pelvic reduction in nine-spined stickleback freshwater populations is universal (see below), such a pattern is consistent with poor connectivity, causing heterogeneity in the patterns of standing genetic variation. This would restrict local adaptation, and hence, also parallel phenotypic evolution. However, the focus of this study is on the role of population demographic parameters on local adaptation from standing genetic variation, regardless of whether the alleles that confer freshwater adaptations are identical by descent (e.g. EDA) or independently derived (e.g. *Pel*-deletions in three-spined sticklebacks), and only secondarily on parallelism (i.e. when the same trait is locally adapted in multiple independently colonised populations, regardless of genetic architecture).

It has been shown that multiple independent alleles for the same adaptation can temporarily co-exist in structured populations, causing heterogeneity in standing genetic variation across the distribution range of a species. This can be true when considering the dynamics of novel mutations (Ralph and Coop 2010) or neutral alleles segregating in the population before they become adaptive (Ralph and Coop 2015a). However, these studies assume uniform selection across single continuous populations and thus do not consider the scenario where locally adapted freshwater alleles – which can be different in different parts of the distribution range – are continually introduced to the sea. Furthermore, simulations showing that the probability of local parallel evolution from standing genetic variation depends on the migration rate from the sea (Galloway and Cresko 2019) or from other populations of the same habitat (Ralph and Coop 2015a) implicitly assume a homogenous pool of ancestral standing genetic variation. Our simulations show that local adaptation is also a function of the allele frequency in the founding marine population; with stronger IBD in the sea, standing genetic variation in the ancestral marine population was a stronger bottleneck for parameters that resulted in nine-spined stickleback-like population structure (9-sp) than for parameters that resulted in three-spined stickleback-like population structure (3-sp; Supplementary Figures 7 and 8). Particularly for smaller effect loci, there was a stronger dependence between the allele frequency in the ancestral marine population and local adaptation in 9-compared to 3-sp, indicating that smaller effect QTL have a stronger influence on local adaptation in 9-compared to 3-sp (Supplementary Figures 7 and 8). This resulted in more polygenic and/or less complete local adaptation in 9-compared to 3-sp (Supplementary Figures 7 and 8), indicating that population demography is sufficient to preclude local adaptation.

While our simulation results match closely with estimates of population structure and IBD in the empirical data, these simulations explore only a small proportion of the possible parameter space with respect to selection intensity, effective population sizes and migration rates (detailed in Supplementary File 4). We also assume that all parallel evolution is due to standing genetic variation of alleles that are identical by descent (since mutation rate is low), whereas it is known that pelvic reduction in three-spined sticklebacks is, to a large extent, due to independently derived *Pel*-deletions attributable to high mutation rates (Xie *et al*. 2019). Nevertheless, most three-spined stickleback studies indicate that freshwater adaptation is indeed chiefly due to alleles identical by descent that have segregated in the population for millions of years (e.g. Jones *et al*. 2012; Nelson and Cresko 2018), making *Pitx1* an exception. In the case of nine-spined sticklebacks, *Pitx1* is with certainty associated with pelvic reduction in only two of all studied populations, suggesting a minor role for recurrent mutation in determining pelvic reduction in this species. While our simulations demonstrate that heterogeneous patterns of standing genetic variation is sufficient for limiting local adaptation in species characterized by low connectivity, further simulations with a wider parameter space could be helpful in advancing our understanding of the limits of adaptation in small and structured populations/species.

Interestingly, in the HEL × BYN cross complete spine reduction most likely occurred when the allele responsible for spine reduction for the LG6 QTL was combined with at least one allele causing spine reduction from the LG15 and LG16 QTL, indicating epistatic interactions (Supplementary File 6). If a threshold number of alleles are needed for complete pelvic reduction, this could also explain how standing genetic variation in the sea is maintained, as the necessary multi-locus genotypes that cause sub-optimal phenotypes in the sea are rarely formed, due to overall lower frequencies of spine-reducing alleles in the sea. This is analogous to “epistatic shielding” that can contribute to the persistence of disease alleles in populations (Phillips and Johnsson 1998; Phillips 2008). Consistent with this possibility, LG6 of the F_0_ female of the HEL × BYN cross (from the sea) was polymorphic for the pelvic spine QTL effect – evidently, a single pelvic-reducing allele alone in this female was not enough to cause any pelvic reduction at all (this female had a complete pelvis).

### Geographic heterogeneity in selection optima

The high heterogeneity in the genetic architecture of pelvic reduction in the nine-spined stickleback can alternatively be attributed to within habitat environmental variation resulting in different selection optima in different pond populations (cf. Stuart *et al*. 2017; Thompson *et al*. 2019). For example, the small differences between pelvic reduced phenotypes in our study (e.g. in BYN and MAS/BÖL spines are completely absent, while in RYT they are only strongly reduced; Fig. 1, Supplementary Table 1 and Supplementary Fig. 3) could in fact indicate different selection optima in the different populations (Stuart *et al*. 2017; Thompson *et al*. 2019). It is possible to use available phenotypic data to estimate the phenotypic optima of pelvic morphology (hypersphere) in each of the populations using Fisher’s geometric model (Stuart *et al*. 2017; Thompson *et al*. 2019), where a strong overlap would suggest a higher probability of genetic parallelism (Thompson *et al*. 2019). However, this assumes that the populations have access to exactly the same ancestral variation and are free to evolve and reach their optima, which is at odds with the results presented here. Without further detailed environmental data or direct estimates of strength of selection on pelvic phenotypes, disentangling the effects of gene flow and within habitat environmental variation (assuming this leads to non-parallel angles of selection) is not possible (Stuart *et al*. 2017).

In a recent simulation study, Thompson *et al*. (2019) showed that genetic parallelism from standing genetic variation rapidly declines as selection changes from fully parallel (optima angle of 0°) to divergent (optima angle of 180°), especially when the trait is polygenic. However, although selection was fully parallel in our simulations, we did not observe strong genetic parallelism for smaller effect loci (with allelic effects < 6) in both the three- and nine-spined stickleback-like scenarios (Fig. 5b, c). This suggests that the effects of the underlying genetic architecture on parallelism (in conjunction with some IBD and population structuring) can be independent of the angle of optimal phenotypes between two habitats. Thus, overlapping selection optima is necessary but nevertheless not sufficient for parallel evolution to take place and it is important not to disregard the population demographic setting as a factor that could severely restrict heritability for adaptation and/or constrain adaptation to less optimal solutions. Evolutionary studies of species with population demographic parameters comparable to those typical for vulnerable or endangered species/populations, such as the nine-spined stickleback, would be valuable to gain a better understanding of how such species may respond to environmental changes and urbanisation (Thompson *et al*. 2018).

### Pelvic reduction outside marine-freshwater study systems

While the evidence for genetic parallelism on large geographical scales in the marine-freshwater stickleback model system is extensive (Colosimo *et al*. 2005; Jones *et al*. 2012; Terekhanova *et al*. 2014, 2019; Nelson and Cresko 2018; Fang *et al*. 2019), the level of parallelism in lake-stream and pelagic-benthic ecotype pairs of three-spined sticklebacks is much more diverse (Peichel *et al*. 2001, 2017; Conte *et al*. 2012, 2015; Stuart *et al*. 2017; Rennison *et al*. 2019). For instance, Conte *et al*. (2015) found that among benthic-limnetic three-spined stickleback pairs from Paxton and Priest lakes (Vancouver Island, BC, Canada), 76% of 42 phenotypic traits diverged in the same direction, whereas only 49% of the underlying QTL evolved in parallel in both lakes. For highly parallel traits in two other pairs of benthic-limnetic sticklebacks, only 32% of the underlying QTL were shared (Conte *et al*. 2012). Similarly, Stuart *et al*. (2017) found that among 11 evolutionary independent replicate pairs of lake-stream three-spined stickleback populations (Vancouver Island, BC, Canada), both within habitat variation and constraints to gene flow contributed to the observed variation in levels of phenotypic parallelism. Different lakes and streams likely do not have similar access to the same global pool of ancestral variation as pairs of marine-freshwater three-spined sticklebacks, where gene flow in the sea is high. This is consistent with the notion that more heterogeneous access to ancestral variation can indeed limit genetic parallelism. This is also true for one example of marine-freshwater three-spined stickleback divergence among isolated insular freshwater populations in the Haida Gwaii archipelago off the northern Pacific coast of Canada (Deagle *et al*. 2013). Here, similar to the nine-spined sticklebacks in this study, several freshwater populations did not display any reduction in pelvic armour. However, those populations that were fully plated were also genetically more similar to adjacent marine individuals, suggesting that recent marine-freshwater admixture and/or selection favouring plated freshwater individuals could explain this pattern. Thus, with respect to access to ancestral variation available for freshwater adaptation, nine-spined sticklebacks are likely closer to the three-spined stickleback lake-stream and benthic-limnetic study systems than to the three-spined stickleback marine-freshwater study system. The only notable exception is the lake-stream three-spined sticklebacks mentioned above, where genetic structuring also is high.

In a small geographic region in Belgium, Raeymaekers et al. (2017) compared the strength and nature of neutral and adaptive divergence along a salinity gradient in nine- and three-spined sticklebacks. Since the lake and pond populations in this study were connected by creaks (allowing contemporary gene flow), in contrast to our study – where a much larger geographic area is considered – population structuring among the nine-spined stickleback population was not stronger than in the three-spined sticklebacks. Despite gene flow, both phenotypic and genetic parallelism was still lower in nine-compared to three-spined sticklebacks. We suspect that lack of parallelism in nine-spined sticklebacks in Raeymaekers et al. (2017) simply lies in lack of ancestral variation – it does not matter how much gene flow there is if the necessary ancestral variation for local adaptation is lacking. Based on our findings here it seems thus likely that ancestral variation for local adaptation in nine-but not three-spined sticklebacks was already lacking in the marine ancestral populations from which the Raeymaekers et al. (2017) study area was colonised from.

### Candidate genes

While the QTL peak for the *Pitx1/Pel* region in the HEL × RYT cross was narrow, this was not the case for the other QTL we detected. Hence, due to the large QTL regions detected by four-way single-mapping analyses, it was not meaningful to perform gene ontology enrichment (GO) analyses – the QTL regions would have contained possibly thousands of genes. Instead, we searched the literature for known candidate genes related to pelvic reduction and found three (excluding *Pitx1*/*Pel*) that were clearly contained within the identified QTL regions (Fig. 2 and Supplementary Table 4). Due to the aforementioned large QTL regions, these can only be considered as highly putative candidate genes for pelvic reduction and will not be discussed further. However, further studies of pelvic reduction might find these candidates worthy of attention.

### Conclusions

Our results show that the repeated parallel reduction in pelvic structures in freshwater populations of nine-spined sticklebacks is due to a diverse set of genetic changes: only one small effect QTL for pelvic reduction was shared between the three experimental crosses in this study. In one cross, pelvic reduction was mapped to the previously identified *Pitx1/Pel* regions, but in the other two crosses, the genetic basis of pelvic reduction was polygenic, and mapped to many different chromosomes. In addition to these, yet another large effect QTL different from the *Pitx1/Pel* locus likely segregates in one nine-spined stickleback population, which is yet to be identified. The results also shed light on the possible drivers of the observed genetic heterogeneity underlying pelvic reduction; as shown by simulations, heterogeneous genetic architectures are more likely to emerge when access to ancestral variation is limited by strong isolation by distance and population structuring. This reinforces the role of the nine-spined sticklebacks as a useful model system, alongside the three-spined stickleback, to study adaptive evolution in the wild. Furthermore, since the population demographic characteristics of nine-spined sticklebacks are similar to small and endangered species/populations, it is also likely to be a well-suited model to study the genetics of adaptation in populations of conservation concern.

## Data availably

The data and code for analyses that support the findings of this study are openly available in Dryad at DOI https://doi.org/10.5061/dryad.76hdr7str. Raw sequence reads have been submitted to NCBIs short read archives with accession numbers: PRJNA673430 and PRJNA672863.

## Acknowledgments

We thank Alexandre Budria, Chris Eberlein, Sami Karja, Heini Natri and Ismo Rautiainen for help in fish collection and/or rearing. Thanks are also due to Kirsi Kahkonen for help in laboratory, and Oulanka Biological Station (University of Oulu) for logistic support. We are grateful for the computing resource support from CSC – the Finnish IT Center for Science Ltd administered by the Ministry of Education and Culture, Finland. B.G. thanks support from CAS Pioneer Hundred Talents Program and the National Natural Science Foundation of China (31672273). This work is supported by the Academy of Finland (grant numbers, 129662, 134728 and 218343 to J.M.), a grant from Helsinki Institute of Life Science (HiLIFE to JM) and a personal grant to Petri Kemppainen from the Finnish Cultural Foundation (00190489).

