## Supplemental documents for "Genetic population structure constrains local adaptation in sticklebacks"

MOLECULAR ECOLOGY

#### Table of Contents:



| | |
|---|---|
| **Supplementary File 1 \| Linkage map construction** | Page 3 |
| **Supplementary File 2 \| Linkage disequilibrium network analyses** | Page 4 |
| **Supplementary File 3 \| Source of QTL effects from four-way single-mapping analyses of $F_2$-intercross data** | Page 5 |
| **Supplementary File 4 \| Simulations** | Page 8 |
| **Supplementary File 5 \| Isolation by distance and population structuring in empirical data** | Page 12 |
| **Supplementary File 6 \| Epistatic interactions** | Page 16 |
| **Supplementary File 7 \| Does local adaptation depend on ancestral allele frequency?** | Page 18 |
| **Supplementary Table 1 \| Summary of phenotypic data** | Page 19 |
| **Supplementary Table 2 \| Sample locations for *Pel*-deletion scan** | Page 20 |
| **Supplementary Table 3 \| Detailed QTL results** | Page 21 |
| **Supplementary Table 4 \| Summary of candidate genes** | Page 24 |
| **Supplementary Figure 1 \| Sample locations for Pel-deletion** | Page 25 |

MOLECULAR ECOLOGY


| | |
|---|---|
| scan | |
| Supplementary Figure 2 \| Outline of simulations | Page 26 |
| Supplementary Figure 3 \| Scatterplots between relative spine and girdle lengths | Page 27 |
| Supplementary Figure 4 \| Fine-mapping of pelvic spine length in LG6, LG15 and LG16 | Page 28 |
| Supplementary Figure 5 \| QTL of pelvic reduction, relative trait values | Page 29 |
| Supplementary Figure 6 \| Sequence alignment showing pel-deletion | Page 30 |
| Supplementary Figure 7 \| Dependency of local adaptation on ancestral variation | Page 31 |
| Supplementary Figure 8 \| QTL-use in parallel Evolution | Page 32 |

#### Supplementary File 1 | Linkage map construction (Lep-MAP3)

Linkage mapping was carried out following the basic LM3 pipeline, and by combining the three crosses. Parental genotypes were called by taking into account the genotype information on offspring, parents and grandparents (module ParentCall2). Markers segregating more distortedly than would be expected by chance (1:1,000 odds) were filtered out (module Filtering2). Loci were then partitioned into chromosomes (modules SeparateChromosomes2 and JoinSingles2) yielding 21 linkage groups with > 75 000 markers assigned to these groups. Finally, the markers were ordered within each linkage group with the module OrderMarkers2, removing markers that were only informative in either the mother or father, respectively. This created two maps for each chromosome, one having more maternal markers and the other having more paternal markers, with an average of ⅔ of the markers shared between the two. Constructing two maps this way removes the effect of markers that are informative only in one parent, as markers informative in different parents are not informative when compared against each other. The phases were converted into grandparental phase by first evaluating the final marker orders and then matching the (parental) phased data with the grandparental data, inverting the parental phases when necessary. Orphan markers from map-ends based on scatter plots of physical and map positions were manually removed.

MOLECULAR ECOLOGY

#### Supplementary File 2 | Linkage disequilibrium network analyses

Initially, all edges (representing LD-values as estimated by $r^2$ using the function snpgdsLDMat from the R-package SNPRelate, Zheng *et al.* 2012) below the LD threshold value of 0.7 were removed. This resulted in many sub-clusters in which all loci were connected by at least a single edge. Second, for each sub-cluster, a complete linkage clustering was performed, where a cluster is defined by its smallest link. Starting from the root, additional sub-clusters were found recursively, where the median LD between all loci was > 0.9. This recursive step was not used in the original implementation, but was later found to increase computational speed and reduce the number of single locus clusters, i.e. result in more efficient complexity reduction. To facilitate computational speed, we only considered SNPs no further than 2000 SNPs away from each other (rather than considering all pairwise values within a chromosome at a time; parameter w2 = 2000), but nevertheless we analysed all SNPs from each cluster at a time (rather than performing LD clustering in windows). Thus, each resulting LD-cluster represents a set of physically adjacent and highly correlated loci. We performed PCA regression based on each LD-cluster in which individuals are separated according to their LD-cluster multi-locus genotypes. Since the loci are highly correlated, most of the genetic variation (>90%) between individuals is explained by the first principal component. For each LD-cluster, we then replaced all the SNP genotypes by the position of the individuals along the first PC axis after removing the sex-linked loci (Pearson's correlation coefficient >0.95 between the PC coordinates and sex). The input data for the LDn-clustering was comprised of the original co-dominant SNP data with individuals from all three crosses pooled. A custom R-code used for this dimensionality reduction is available from DRYAD (DOI https://doi.org/10.5061/dryad.76hdr7str).

MOLECULAR ECOLOGY

**Supplementary File 3 | Source of QTL effects from four-way single-mapping analyses of $F_2$-intercross data.**

Here we explain in more detail what information about the source of the QTL effects can be inferred from four-way single-mapping analyses when the parental phase is known. Consider the four causal alleles *A*, *B*, *C* and *D*. For the $F_0$ generation we start with crossing the following two genotypes:

$$F_0 : AB \times CD$$

From this cross, four distinct genotypes in the $F_1$ generation can be produced; *AC* (1), *AD* (2), *BC* (3) and *BD* (4). Since only two $F_1$ individuals are chosen to produce the $F_2$ generation, the outcome for the QTL cross will depend on exactly which two of the four possible $F_1$ parental genotypes were sampled (10 possible combinations). For instance, if the $F_1$ genotypes are as follows (genotypes 1 and 4):

$$F_1 : AC(1) \times BD(4)$$

the coding systems derived from bi-allelic SNP data for the $F_2$ offspring [$X_{dij}$, $X_{sij}$, $Z_{ij}$] can be specified as follows:

$$\begin{vmatrix} +1 +1 +1 \\ +1 -1 -1 \\ -1 +1 -1 \\ -1 -1 +1 \end{vmatrix} \begin{matrix} forgenotypeAC, \\ forAD, \\ forBC, \\ forBD. \end{matrix}$$

Here it can be noted the alleles *A* and *B* always come from the $F_1$ male (left, i.e. the grandfather) and alleles *C* and *D* from the $F_1$ female (right, i.e. the grandmother). When the allelic effects of allele *A* = *B* and those of *C* = *D* and alleles *A* and *B* are recessive (relative to *C* and *D*), the following phenotypic values can be expected:

$$\begin{vmatrix} 1 \\ -1 \\ -1 \\ -1 \end{vmatrix} \begin{matrix} forgenotypeAC, \\ forAD, \\ forBC, \\ forBD. \end{matrix}$$

Here a significant QTL can be expected for all three coding systems $X_{dij}$, $X_{sij}$, $Z_{ij}$, as is the case for the *Pitx*1 QTL on LG7 in the HEL × RYT cross (Fig 2 a-b, main text). Note that even if a significant QTL for the coding system $Z_{ij}$ will indicate dominance, further analyses are needed to infer which sets of the grandparental alleles (*AB* versus *CD*) are recessive/dominant.

If instead the sets of the grandparental alleles (*AB* versus *CD*) are additive and the allelic effects of *A* = *B* and those of *C* = *D*, the following phenotypes can be expected:

$$\left| \begin{array}{l} 1 \\ 0 \\ 0 \\ -1 \end{array} \right| \begin{array}{l} for genotype AC, \\ for AD, \\ for BC, \\ for BD. \end{array}$$

and significant QTL can be expected for coding systems $X_{dij}$, $X_{sij}$ but not $Z_{ij}$. Thus, in these simple cases, no additional information about the QTL effects is gained relative to analyses that do not take the grandparental phase into account. However, it is also possible that the allelic effects of *A* ≠ *B*. In this case, several potential outcomes are possible (still assuming the allelic effects of C = *D*). If for instance the $F_1$ genotypes are as follows:

$$F_1 : AC(1) \times BD(4)$$

The following coding system from bi-allelic SNP data can be specified for the $F_2$ offspring [$X_{dij}$, $X_{sij}$, $Z_{ij}$]:

$$\left| \begin{array}{l} +1+1+1 \\ +1-1-1 \\ -1+1-1 \\ -1-1+1 \end{array} \right| \begin{array}{l} for genotype AA, \\ for AD, \\ for CA, \\ for CD. \end{array}$$

Since here the *B* allele is not sampled (i.e. not present among the two $F_1$ parents), the following phenotypes for the $F_2$ offspring can be expected:

$$\left| \begin{array}{l} 1 \\ 0 \\ 0 \\ -1 \end{array} \right| \begin{array}{l} for genotype AA, \\ for AD, \\ for CA, \\ for CD. \end{array}$$

and the expected outcome for the QTL effects is the same as when the allelic effect of *A* = *B* (see above). However, if instead the $F_1$ genotypes are as follows:

$$F_1 : AC(1) \times BD(4)$$

and the additive allelic effect for *A* is 1 and -1 for the remaining alleles (i.e. the allelic effects of *C* = *D*), the following phenotypes for the $F_2$ offspring can be expected:

MOLECULAR ECOLOGY

$$\left| \begin{array}{l|l} 1 & for genotype AC, \\ 1 & for AD, \\ -1 & for BC, \\ -1 & for BD. \end{array} \right.$$

In this scenario, a significant QTL can be expected only for the coding system $X_{dij}$. From this, we can infer that *i*) the grandfather was heterozygous for different causal variants (i.e. the allelic effects of *A* ≠ *B*) and *ii*) the two $F_1$ parents must have inherited different grandpaternal causal alleles (*A* and *B*). This is the case for QTL for spine length on LG15 and LG16 for the HEL × BYN cross (Fig 2 b-e, main text). If instead (in the above cross) the allelic effect of *C* is 1 and that of the remaining alleles is -1, a significant QTL can be expected only for the coding system $X_{sij}$, and thus the allelic effects of C ≠ *D* (i.e. the SNP inherited from the grandmother must have been linked (on different chromatids) to the different causal alleles *C* and *D*).

Thus, while additional information about the allelic QTL effects can be directly gained using four-way single-mapping analyses, this ultimately also depends on exactly which of the four potential genotypes from the $F_1$ generation were sampled to produce the $F_2$ generation. Note also that since the causal alleles *A and B* (from the grandfather) and *C and D* (from the grandmother) can all have different allelic effects and dominance relationships, the interpretation of the outcome from the four-way single-mapping analyses can potentially become highly complex and complicated to interpret.

When producing the coding systems $X_{dij}$, $X_{sij}$, $Z_{ij}$, it is possible for one of the $F_0$ SNP genotypes to be heterozygous as long as both of the $F_1$ individuals are heterozygous. However, from the $F_0$ individual that is heterozygous, it is of course not possible to infer different allelic effects for the two different grandparental phases.

MOLECULAR ECOLOGY

#### Supplementary File 4 | Simulations

For both nine- and three-spined stickleback-like population demographic scenarios (henceforth 9- and 3-sp, respectively), the sea populations were simulated by a stepping-stone model comprised of ten sub-populations, each with carrying capacity K = 1000 (under mutation-drift equilibrium K = $N_e$). All adjacent marine populations exchanged 100 (3-sp) or ten (9-sp) migrants per generation (M, symmetrical gene flow), with some long-distance migration also allowed between every second population at a rate of ten times less than that between adjacent populations (Supplementary Fig. 2). To allow frequencies of freshwater adapted alleles to build up and filter to the marine populations as standing genetic variation, a refuge freshwater population (K = 10000) with high frequencies of freshwater adapted alleles was allowed to exchange migrants with the two most central of the ten marine populations (symmetric gene flow with rate M = 1; Supplementary Fig. 2). After a burn-in, two focal freshwater populations were founded from the two marine populations situated at opposite ends of the stepping-stone chain, after which symmetrical gene flow was allowed between the focal freshwater populations and their closest marine populations at rates M = 1 for 3-sp and M = 0.2 for 9-sp (Supplementary Fig. 3). Thus, the freshwater populations were founded from different marine populations with ancestral freshwater adapted alleles stemming from the same refuge freshwater population situated equally far from the two focal freshwater populations, in agreement with the transporter hypothesis. The simulations were then run for 5000 generations to simulate post glacial colonisation of freshwater habitats (10000 years ago, assuming a generation time of two; Baker 1994; DeFaveri and Merilä 2013; DeFaveri *et al.* 2014). The above parameters generated patterns of IBD and population structuring similar to what was estimated from the empirical data (see Results), and are also consistent with the results of previous studies (DeFaveri *et al.* 2012).

Three different genetic architectures of a single trait coded by five (independent) bi-allelic loci under stabilising selection for different optima were simulated in the marine (optimal phenotypic value = 0) and freshwater populations (optimal phenotypic value = 20). In architecture A, one large effect additive locus with the homozygote for allele 1 ("a1") yielding a genotypic value of zero (locally adapted to the marine habitat) was simulated; the same was done for the homozygote for allele 2 ("a2") yielding a genotypic value of 20 (locally adapted for the freshwater habitat). The four remaining minor effect additive QTL were given the allelic value of zero for a1 and allelic values [1,1,2,3] for a2. An empirical example of this kind of architecture is the Ectodysplasin (*EDA*) gene in the 3-sp (Colosimo *et al.* 2005). In architecture B, the allelic values of a2 alleles were [1,2,3,4,6] and the allelic values of the alternative alleles were zero, i.e. a single large effect locus is lacking. In

MOLECULAR ECOLOGY

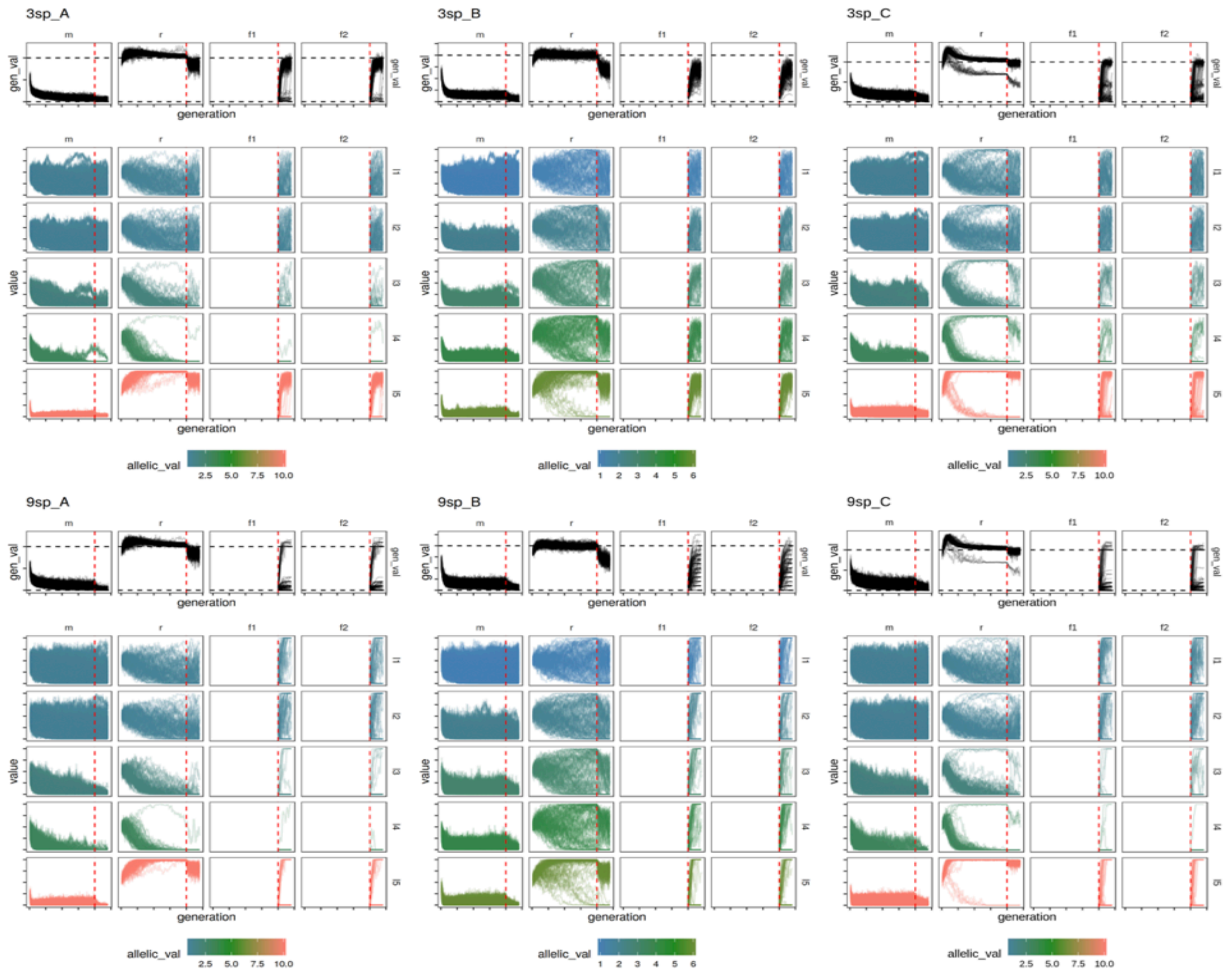


**Allele frequency trajectories and genotypic values for simulations.** Allele frequency trajectories (bottom) for all QTL are shown throughout the simulations separately for the refugee ("r"), the two freshwater populations ("f1" and "f2"), and all marine populations (pooled; "m2). Species (3sp for three-spined and 9sp for nine-spined sticklebacks), and genetic architecture (A, B or C) is indicated by the title for each sub-graph. Allele frequencies start at 0.5 and the range for the y-axis is zero to one. Colour indicates allelic effect sizes for locus 1 (l1) to locus 5 (l5). Top panel shows trajectories for genotypic values (based on all QTL), with the top horizontal line indicating 20 (i.e. the optimal genotypic value for freshwater), and the bottom horizontal line indicating zero (i.e. the optimal genotypic value for local adaptation to the sea. Red vertical line marks the end of burn-in and colonisation of the two freshwater populations.

architecture C, the distribution of allelic values was the same as in architecture A, but with a2 being recessive to a1. An empirical example of this is the *Pitx1* locus controlling pelvic reduction in 3- and 9-sp (Cresko *et al.* 2004; Shapiro *et al.* 2004). All traits were treated as quantitative and their fitness (*W*) was mapped to the phenotypes (*P*) using a standard Gaussian function: $W$=e^(-($p$-$z_h$)/2$s^2$), where $z_h$ is the phenotypic optima in habitat *h* and the selection intensity (*s*) was set to 100 for freshwater habitats and 200 for the marine habitats, with lower values translating to stronger selection intensity (see quantiNemo manual for details). Lower selection intensity in the sea allowed higher frequencies of the freshwater allele (allele 2). This allowed for rapid local adaptation (Supplementary Figure

6) in the freshwater habitats following colonisation from the sea, conditional on that there was sufficient standing genetic variation in the ancestral sea population (figure above).

We simulated ten chromosomes of 100 centi Morgan (cM) with 100 neutral loci (and the QTL) equally spaced across the genome (i.e. uniform recombination rate across chromosomes, with no tight linkage between any two loci). Mutation rate (μ) was set to $1e^{-8}$ for all loci, and simulations were initiated with allele frequencies for all loci set to 0.5. At the end of the simulations (generation 15000) we estimated allele frequency differences as the fixation index, $F_{ST}$ (Weir and Cocherham 1984), with the stamppFst function in the R-package StAMPP (Pembleton *et al.* 2013). Further details of parameter settings and the simulation code can be found from DRYAD (DOI https://doi.org/10.5061/dryad.76hdr7str).

We classified loci being involved in parallel evolution if the $F_{ST}$ between the focal freshwater populations (pooled) and all marine populations (pooled) was higher than 0.5 (marine-freshwater $F_{ST}$) and lower than 0.5 between the two focal freshwater populations (freshwater-freshwater $F_{ST}$). Any loci with both high marine-freshwater $F_{ST}$ and freshwater-freshwater $F_{ST}$ (above 0.5) were classified as being locally adapted, but not involved in parallel evolution.

MOLECULAR ECOLOGY

#### Supplementary File 5 | Empirical data set

*Sampling, sequencing and genotype calling*

We obtained sequence datasets of three-spined sticklebacks from Fang *et al.* (2020) and nine-spined sticklebacks from Fang *et al.* (2020). The European sticklebacks from these studies are the focal populations for the analyses of isolation by distance (IBD), since their sampling covers populations from both marine and freshwater habitats in both species. To maximally cover comparable samples from both species, the available whole genome sequencing (WGS) and restriction site associated DNA sequencing (RADseq) datasets were included, and only the populations with at least two individuals were included. In total, 25 three-spined stickleback populations (16 freshwater and 9 marine) and 31 nine-spined stickleback populations (14 freshwater and 7 marine) were used for IBD analyses. See table below for sampling details.

In each species, all sequence datasets were mapped to the reference genome of the corresponding species. The reference genome of the three-spined stickleback (release-92) was retrieved from Ensembl (Hubbard *et al.* 2005), and that of the nine-spined stickleback was retrieved from a recent assembled reference genome (Varadharajan *et al.* 2019). The genotyping was performed on mapped reads (in BAM format) using BWA mem v0.7.17 (Li and Durbin, 2010), SAMtools packages v1.4 (Li, 2011) and Genome Analysis Toolkit (GATK v3.7; DePristo *et al.* 2011). Downstream filtering steps were performed with VCFtools v. 0.1.15 (Danecek *et al.* 2011). Sex chromosomes in each species were excluded (chromosome XIX for three-spined stickleback; Kitano *et al.* 2009; Natri *et al.* 2013; LG 12 for nine-spined stickleback; Shapiro *et al*. 2009; Rastas *et al.* 2016). We retained only bi-allelic SNPs with less than 20% missing genotypes (--max-missing 0.8, --min-alleles 2 --max-alleles 2), minor allele frequency above 0.02 (--maf 0.02), and heterozygosity below 0.5 using, a python script (github.com/z0on/2bRAD_GATK) to remove potential paralogs, and thinned the final dataset (--thin 30,000) so that no loci were closer than 30 kb from each other. The raw output from the individuals comprised 4,701,869 loci for three-spined and 12,709,617 for nine-spined sticklebacks (allowing 25% missing genotypes initially); this was filtered to a set of 10,174 SNPs for three-spined and 4,326 SNPs for nine-spined sticklebacks. To match the dataset sizes, 4,326 SNPs were randomly selected from the three-spined stickleback dataset for the analyses. These data will be available from NCBIs short read archives via accession number: PRJNA672863. As the data here is a small subset of a much larger data set used for an unpublished stickleback comparative study, the raw sequences are currently based under embargo.

### Table. Sample summary of the empirical data set

| Species | Population ID | Ecotype | Region | GPS_N | GPS_E | Sample size | Source of sequences |
|---|---|---|---|---|---|---|---|
| Three-spined stickleback | BOL | Freshwater | White and Barents Sea | 66.295278 | 33.366111 | 10 | Fang et al. (2019) |
| Three-spined stickleback | MAS | Freshwater | White and Barents Sea | 66.291944 | 33.381667 | 2 | Fang et al. (2019) |
| Three-spined stickleback | KVA | Freshwater | White and Barents Sea | 70.094444 | 28.978611 | 10 | Fang et al. (2019) |
| Three-spined stickleback | KEV | Freshwater | White and Barents Sea | 69.750278 | 27.015278 | 9 | Fang et al. (2019) |
| Three-spined stickleback | ROT | Freshwater | Baltic Sea | 64.265000 | 20.318889 | 2 | Fang et al. (2019) |
| Three-spined stickleback | VAT | Freshwater | Baltic Sea | 58.646111 | 14.638611 | 2 | Fang et al. (2019) |
| Three-spined stickleback | SLI | Freshwater | Baltic Sea | 59.936944 | 31.022778 | 10 | Fang et al. (2019) |
| Three-spined stickleback | NEV | Freshwater | Baltic Sea | 54.638889 | 25.364167 | 2 | Fang et al. (2019) |
| Three-spined stickleback | SKF | Freshwater | Norwegian Sea | 69.956667 | 19.145833 | 9 | Fang et al. (2019) |
| Three-spined stickleback | TAK | Freshwater | Norwegian Sea | 69.117222 | 19.067500 | 5 | Fang et al. (2019) |
| Three-spined stickleback | FAR | Freshwater | Norwegian Sea | 62.150278 | -6.633889 | 2 | Fang et al. (2019) |
| Three-spined stickleback | MYV | Freshwater | Norwegian Sea | 65.642222 | -16.927222 | 5 | Fang et al. (2019) |
| Three-spined stickleback | MYR | Freshwater | North Sea and British Isles | 60.313333 | 5.399167 | 10 | Fang et al. (2019) |
| Three-spined stickleback | KIN | Freshwater | North Sea and British Isles | 56.335000 | -2.789722 | 2 | Fang et al. (2019) |
| Three-spined stickleback | QUI | Freshwater | North Sea and British Isles | 55.789167 | -5.088611 | 2 | Fang et al. (2019) |
| Three-spined stickleback | BUT | Freshwater | North Sea and British Isles | 51.728611 | 0.160556 | 2 | Fang et al. (2019) |
| Three-spined stickleback | PRI | Marine | Baltic Sea | 60.350000 | 28.616667 | 2 | Fang et al. (2019) |
| Three-spined stickleback | BAR | Marine | White and Barents Sea | 74.966944 | 37.134167 | 2 | Fang et al. (2019) |
| Three-spined stickleback | SBJ | Marine | White and Barents Sea | 71.823333 | 33.043333 | 2 | Fang et al. (2019) |
| Three-spined stickleback | IND | Marine | White and Barents Sea | 66.240000 | 37.145000 | 2 | Fang et al. (2019) |
| Three-spined stickleback | LEV | Marine | White and Barents Sea | 66.290556 | 33.434167 | 2 | Fang et al. (2019) |
| Three-spined stickleback | KRI | Marine | North Sea and British Isles | 58.165556 | 8.031111 | 2 | Fang et al. (2019) |
| Three-spined stickleback | FIS | Marine | North Sea and British Isles | 58.234722 | 11.401667 | 2 | Fang et al. (2019) |
| Three-spined stickleback | BS_3sp | Marine | North Sea and British Isles | 56.602553 | 8.300539 | 6 | Fang et al. (2019) |
| Three-spined stickleback | Ran_M | Marine | North Sea and British Isles | 56.606686 | 10.302019 | 3 | Fang et al. (2019) |
| Nine-spined stickleback | RUS-BOL | Freshwater | White and Barents Sea | 66.3 | 33.4 | 6 | Fang et al. (2020) |
| Nine-spined stickleback | RUS-MAS | Freshwater | White and Barents Sea | 66.3 | 33.4 | 6 | Fang et al. (2020) |
| Nine-spined stickleback | RUS-KRU | Freshwater | White and Barents Sea | 66.3 | 33.4 | 6 | Fang et al. (2020) |
| Nine-spined stickleback | FIN-PUL | Freshwater | White and Barents Sea | 69.96666667 | 27.9666 | 6 | Fang et al. (2020) |
| Nine-spined stickleback | FIN-KEV | Freshwater | White and Barents Sea | 69.75 | 27.01666667 | 6 | Fang et al. (2020) |
| Nine-spined stickleback | VEN-AAN | Freshwater | White and Barents Sea | 61.58333333 | 34.63333333 | 2 | Fang et al. (2020) |
| Nine-spined stickleback | SWE-ABB | Freshwater | Baltic Sea | 64.47833333 | 19.43638889 | 4 | Fang et al. (2020) |
| Nine-spined stickleback | SWE-BYN | Freshwater | Baltic Sea | 64.45527778 | 19.44444444 | 3 | Fang et al. (2020) |
| Nine-spined stickleback | SWE-NAV | Freshwater | Baltic Sea | 64.56472222 | 19.19861111 | 3 | Fang et al. (2020) |
| Nine-spined stickleback | SWE-HAN | Freshwater | Baltic Sea | 64.55666667 | 19.17388889 | 3 | Fang et al. (2020) |
| Nine-spined stickleback | DEN-RES | Freshwater | North Sea and British Isles | 56.183333 | 9.633333 | 3 | Fang et al. (2020) |
| Nine-spined stickleback | PP-FIN-PAL | Freshwater | White and Barents Sea | 68.02861111 | 24.15722222 | 6 | Fang et al. (2020) |
| Nine-spined stickleback | PP-NO-STO | Freshwater | Norwegian Sea | 69.747778 | 18.415000 | 3 | Fang et al. (2020) |
| Nine-spined stickleback | NOR-UGE | Freshwater | Norwegian Sea | 63.9584775 | 10.42684639 | 20 | Fang et al. (2020) |
| Nine-spined stickleback | PP-FR-MAR | Freshwater | North Sea and British Isles | 47.024167 | 5.254444 | 2 | Fang et al. (2020) |
| Nine-spined stickleback | BEL-MAL | Freshwater | North Sea and British Isles | 51.174722 | 3.469444 | 5 | Fang et al. (2020) |
| Nine-spined stickleback | PP-UK-HAR | Freshwater | North Sea and British Isles | 55.753889 | -4.428889 | 2 | Fang et al. (2020) |
| Nine-spined stickleback | PP-UK-LRE | Freshwater | North Sea and British Isles | 57.61103889 | -7.514886111 | 2 | Fang et al. (2020) |
| Nine-spined stickleback | PP-UK-LSC | Freshwater | North Sea and British Isles | 57.58436111 | -7.236211111 | 2 | Fang et al. (2020) |
| Nine-spined stickleback | GBR-GRO | Freshwater | North Sea and British Isles | 57.615122 | -7.511461 | 6 | Fang et al. (2020) |
| Nine-spined stickleback | FIN-KAR | Freshwater | Baltic Sea/White Sea | 66.437222 | 29.135556 | 20 | Fang et al. (2020) |
| Nine-spined stickleback | FIN-KRK | Freshwater | Baltic Sea/White Sea | 66.656667 | 26.440833 | 20 | Fang et al. (2020) |
| Nine-spined stickleback | FIN-PYO | Freshwater | Baltic Sea/White Sea | 66.261111 | 29.433333 | 31 | Fang et al. (2020) |
| Nine-spined stickleback | FIN-RYT | Freshwater | Baltic Sea/White Sea | 66.384167 | 29.320000 | 21 | Fang et al. (2020) |

| | | | | | | | |
|---|---|---|---|---|---|---|---|
| Nine-spined stickleback | GER-RUE | Marine | Baltic Sea | 54.401506 | 13.210098 | 6 | Fang et al. (2020) |
| Nine-spined stickleback | FI-HEL | Marine | Baltic Sea | 60.202507 | 25.182756 | 6 | Fang et al. (2020) |
| Nine-spined stickleback | SWE-BOL | Marine | Baltic Sea | 63.661389 | 20.211756 | 6 | Fang et al. (2020) |
| Nine-spined stickleback | FIN-KIV | Marine | Baltic Sea | 65.007945 | 25.435659 | 6 | Fang et al. (2020) |
| Nine-spined stickleback | RUS-LEV | Marine | White and Barents Sea | 66.373610 | 33.768502 | 6 | Fang et al. (2020) |
| Nine-spined stickleback | DEN-NOR | Marine | North Sea and British Isles | 54.985148 | 8.659472 | 9 | Fang et al. (2020) |
| Nine-spined stickleback | SWE-FIS | Marine | North Sea and British Isles | 58.23333333 | 11.4 | 9 | Fang et al. (2020) |

MOLECULAR ECOLOGY

#### Supplementary File 6 | Epistatic interactions

For complex traits, epistasis describes any interaction between two or more loci, such that the phenotype of any genotype cannot be predicted simply by summing the effects of the individual loci (Carlborg and Haley 2004). Thus, in the absence of any large effect loci, and when all QTL are additive and independent, as is the case for the QTL on LGs 6, 15 and 16 for the HEL × RYT cross, phenotypes in the F2 generation are expected to be normally distributed (Klug and Cummings 2018). For some multi-locus genotypes (of the QTL on LGs 6, 15 and 16), the distribution of spine lengths (figure below) was approximately normally distributed, except the highly reduced spine lengths, which had long tails, implicating that some of these loci could be involved in epistatic interactions.

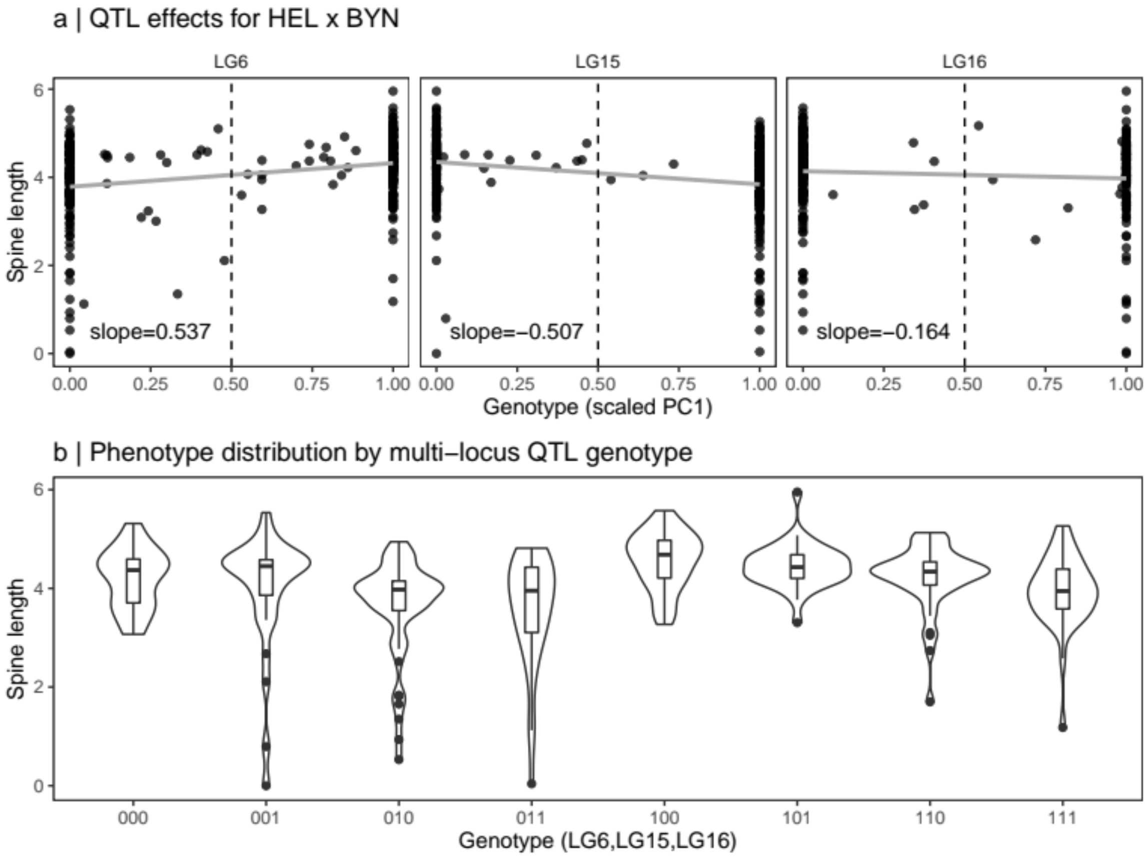


**Epistatic interactions in pelvic spine development for HEL × BYN cross.** (a) The effects of individual genotype, where the genotype is given by the first PC (scaled between 0 and 1) from the cluster of SNPs that was the most significant for spine length on LG6, LG15 and LG16 (see Fig. 2, main text), respectively. Genotypes were further based on the genotypes [$x_{dij}$, $x_{sij}$] depending on which of these were significant for the QTL effects ($x_{dij}$ for LG6 and $x_{sij}$ for LG15 and LG16). Some individuals have genotypes between 0 and 1 only because the genotype is based on the first PC of large sets of highly but not perfectly correlated SNPs. Slope of the regression line (grey) is shown. (b) distribution of spine lengths for all multi-locus genotypes from (a). The multi-locus genotype was based on rounding PC1 coordinates from (a) where values below 0.5 (left of vertical dashed line) were considered as "Allele 0", and those above or equal to 0.5 were considered as "Allele 1". The first digit for genotypes in (b) thus represents the genotype of the QTL on LG6, followed by LG15 and LG16, respectively.

MOLECULAR ECOLOGY

This possibility was investigated as follows. Individual allelic effects were visualised (figure above) by finding LD-clusters with the most significant QTL effect (using nominal P-values from the four-way single-mapping analyses) and regressing the coordinates from PC1 (based on the loci from these sets of highly correlated loci), scaled to values [0,1], against relative spine length. Only the relevant coding systems shown in equation (2) in the main text were used, for which the QTL effect was significant. When testing for possible interactions, we rounded the scaled PC1 coordinates to the closest zero or one, thus retaining a binary genotype (alleles “0” and “1”) for each of the QTL and the coding system. In particular, we observe that the multi-locus genotypes “011”, “010” and “001” have high proportions of the most reduced phenotypes, compared to the remaining multi-locus genotypes. We therefore tested whether among $n_l$-th smallest phenotypic values we observed more “011”, “010” and “001” genotypes (pooled) than from the remaining genotypes (pooled) than expected under the null hypothesis of no association between multi-locus genotypes and phenotype, using Fishers exact tests for $n_l$[10,50]. Due to the limited numbers of individuals with fully reduced spines, it was not possible to perform more elaborate analyses such as epistatic QTL mapping (Carlborg and Haley 2004). Results are given in the figure caption below.

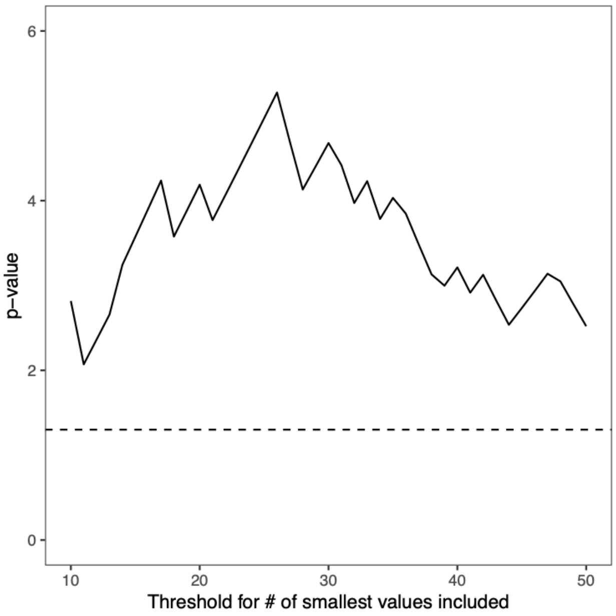


Figure to the left shows -log10 p-values (y-axis) for Fishers exact test as a function of how many of the smallest relative spine lengths were included in the test (x-axis). The test remained significant (points above the horizontal line that represents significance at the 5% level) for the whole range of $n_l$ values tested [10,50]. Thus, the large proportion of the smaller phenotypic values observed for the multi-locus genotypes “011”, “010” and “001” is unlikely to happen by chance.

MOLECULAR ECOLOGY

**Supplementary File 7 | Does local adaptation depend on ancestral allele frequency?**

In the simulations, the frequency of a2 allele (locally adapted to freshwater) in each freshwater population after 2000 generations post-colonisation from the sea was always correlated with the a2 allele frequency in the founding marine population, demonstrating that local adaptation depended on ancestral genetic variation (Supplementary Fig. 7). This dependency was the strongest for small effect loci (phenotypic effect of a2 = 1). In these cases, the slope (β) of a linear regression between the a2 frequency in the sea and in the freshwater population, respectively, ranged between 1 and 1.4, the y-intercept ranged between 0 and 0.09, and Pearson's squared correlation coefficient ($r^2$) ranged between 0.33 and 0.52, and was lowest with parameter settings resulting in three-spined stickleback-like population structures (hereafter 3-sp) for genetic architecture A (y-intercept = 0.51; β = 4.2; $r^2$ = 0.07; Supplementary Fig. 7). In the latter case, standing genetic variation (defined as a2 frequency > 0) was available in 89% of the simulation replicates (considering each freshwater population independently), of which 76% resulted in local adaptation (defined as a2 frequency > 0.5) in at least one of the freshwater populations. The corresponding numbers for nine-spined stickleback-like population structures (hereafter 9-sp) were 41% and 39%, respectively. Here the correlation between ancestral a2 frequency and a2 frequency post-colonisation was also stronger for 9- than 3-sp (y-intercept = 0.04; β = 10.8; $r^2$ = 0.33), indicating a stronger dependence between ancestral variation and local adaptation. When standing genetic variation was available for local adaptation, three factors likely contributed to lower levels of local adaptation and parallel evolution in 9- relative to 3-sp : i) smaller K freshwater populations (500 *vs.* 1000), which is expected to result in faster genetic drift; ii) lower (1/5th ) post-colonisation gene-flow between adjacent marine and freshwater populations; and iii) lower a2 frequency (when present as standing genetic variation) in 9- compared to 3-sp sticklebacks (2.9% vs. 3.4%). Thus, IBD in the sea likely limits the probability that a freshwater adapted allele is maintained as standing genetic variation, while smaller effective population sizes and reduced gene flow can further limit local adaptation, even when ancestral variation in the founding population is present (in low frequencies). The generally higher β's (3.20 *vs*. 2.15) and y-intercepts (0.11 *vs*. 0.05) for QTL with a2 < 6 for 9-sp indicate that smaller effect QTL mostly have a stronger influence on local adaptation in 9- compared to 3-sp (Supplementary Fig. 7), resulting in polygenic and/or incomplete local adaptation (Supplementary Fig. 7 and Supplementary Fig. 8).

MOLECULAR ECOLOGY

**Supplementary Table 1 | Summary of phenotypic data.** “CG” indicates that data is from common garden experiments. Mean and standard deviations (SD) are based on relative trait values. $r^2$ and *P*-values refer to squared correlation and P-values from regressions of standardised girdle on spine lengths in given population, respectively.

| Population | Category | Mean spine | SD spine | Mean girdle | SD girdle | $r^2_{spine\text{-}girdle}$ | $P_{spine\text{-}girdle}$ |
|---|---|---|---|---|---|---|---|
| **BÖL** | marine | 0.11 | 0.012 | 0.165 | 0.009 | 0.01 | 0.72 |
| **FIS** | marine | 0.105 | 0.012 | 0.184 | 0.011 | 0.00 | 0.98 |
| **HEL** | marine | 0.115 | 0.012 | 0.172 | 0.008 | 0.01 | 0.64 |
| **LEV** | marine | 0.107 | 0.01 | 0.17 | 0.011 | 0.10 | 0.09 |
| **TRE** | marine | 0.117 | 0.013 | 0.157 | 0.01 | 0.29 | 0.01 |
| **HEL_CG** | marine (CG) | 0.098 | 0.008 | 0.145 | 0.013 | 0.02 | 0.42 |
| **LEV_CG** | marine (CG) | 0.088 | 0.005 | 0.137 | 0.006 | 0.19 | 0.01 |
| **JOR** | lake | 0.094 | 0.01 | 0.151 | 0.012 | 0.00 | 0.85 |
| **POR** | lake | 0.088 | 0.009 | 0.143 | 0.006 | 0.08 | 0.14 |
| **RII** | lake | 0.088 | 0.006 | 0.171 | 0.01 | 0.14 | 0.09 |
| **SKA** | lake | 0.081 | 0.008 | 0.132 | 0.006 | 0.12 | 0.06 |
| **ABB** | pond | 0.055 | 0.01 | 0.137 | 0.01 | 0.04 | 0.36 |
| **BOL** | pond | 0.051 | 0.037 | 0.125 | 0.047 | 0.61 | <0.01 |
| **BYN** | pond | 0 | 0 | 0.12 | 0.014 | na | na |
| **HAN** | pond | 0.073 | 0.018 | 0.138 | 0.009 | 0.00 | 0.9 |
| **KAR** | pond | 0.118 | 0.008 | 0.186 | 0.014 | 0.01 | 0.75 |
| **KRK** | pond | 0.073 | 0.016 | 0.151 | 0.007 | 0.01 | 0.69 |
| **MAS** | pond | 0 | 0 | 0 | 0 | na | na |
| **NAV** | pond | 0.081 | 0.007 | 0.149 | 0.007 | 0.00 | 0.81 |
| **PYÖ** | pond | 0.072 | 0.007 | 0.144 | 0.008 | 0.19 | 0.03 |
| **RYT** | pond | 0 | 0 | 0.087 | 0.017 | na | na |
| **BYN_CG** | pond (CG) | 0.033 | 0.018 | 0.1 | 0.01 | 0.03 | 0.3 |
| **PYÖ_CG** | pond (CG) | 0.064 | 0.003 | 0.116 | 0.007 | 0.06 | 0.17 |
| **HEL × RYT** | F2 | 0.055 | 0.033 | 0.138 | 0.041 | 0.85 | <0.01 |
| **HEL × BYN** | F2 | 0.078 | 0.017 | 0.176 | 0.012 | 0.11 | <0.01 |
| **HEL × PYÖ** | F2 | 0.091 | 0.009 | 0.169 | 0.012 | 0.08 | <0.01 |

MOLECULAR ECOLOGY

**Supplementary Table 2 | Sample locations for Pel-deletion scan**

| Name | Lat | long | type |
|---|---|---|---|
| **BEL-MAL** | 51°10'N | 03°28'E | Freshwater |
| **DEN-NOR** | 55°59'N | 08°39'E | Marine |
| **DEN-RES** | 56°11'N | 9°38'E | Freshwater |
| **FIN-HEL** | 60°13'N | 25°11'E | Marine |
| **FIN-KAR** | 66°39'N | 26°26'E | Pond |
| **FIN-KIV** | 65°00'N | 25°28'E | Marine |
| **FIN-KRK** | 66°26'N | 29°08'E | Pond |
| **FIN-PUL** | 69°58'N | 27°58'E | Lake |
| **FIN-PYO** | 66°15'N | 29°26'E | Pond |
| **FIN-RYT** | 66°23'N | 29°19'E | Pond |
| **GBR-GRO** | 57°37'N | -7°30'E | Freshwater |
| **GER-RUE** | 54°24'N | 13°12'E | Marine |
| **NOR-UGE** | 63°57'N | 10°24'E | Freshwater |
| **RUS-BOL** | 66°18'N | 33°24'E | Pond |
| **RUS-KRU** | 66°18'N | 33°25'E | Freshwater |
| **RUS-LEN** | 72°23'N | 126°30'E | Freshwater |
| **RUS-LEV** | 66°18'N | 33°25'E | Marine |
| **RUS-MAS** | 66°18'N | 33°25'E | Pond |
| **SWE-ABB** | 64°29'N | 19°26'E | Pond |
| **SWE-BOL** | 63°39'N | 20°12'E | Marine |
| **SWE-BYN** | 64°27'N | 19°26'E | Pond |
| **SWE-KIR** | 67°53'N | 20°6'E | Freshwater |
| **SWE-FIS** | 58°14'N | 11°24'E | Marine |
| **SWE-HAN** | 64°33'N | 19°10'E | Pond |
| **SWE-NAV** | 64°34'N | 19°12'E | Pond |
| **USA-HLA** | 61°35'N | -149°45'E | Freshwater |

MOLECULAR ECOLOGY

**Supplementary Table 3 | Detailed QTL results.** Results for each individual LD-cluster of highly correlated SNPs. "Pos" refers to position in cM; "Std." refers to whether a trait value was standardised or not; "*P*" and "$P_{cor}$" refer to nominal and corrected *P*-values, respectively; "PVE" refers to the proportion of phenotypic variance explained by the cluster; "$PVE_{TOT}$" refers to the total amount of PVE explained by the peak (all LD-clusters from a given QTL region considered jointly) and "β" is the effect size. Part 1/3.

| Cross | Trait | Coding | LG | QTL | Pos | Std. | *P* | $P_{COR}$ | PVE | $PVE_{TOT}$ | *β* |
|---|---|---|---|---|---|---|---|---|---|---|---|
| **HEL × RYT** | Spine | Male | 7 | 1 | 52.8 | No | 1.31e-04 | 0.019 | 0.056 | 0.285 | 0.673 |
| **HEL × RYT** | Spine | Female | 7 | 1 | 52.8 | No | 1.26e-04 | 0.026 | 0.057 | 0.311 | 0.678 |
| **HEL × RYT** | Girdle | Male | 7 | 1 | 52.8 | No | 3.66e-04 | 0.04 | 0.048 | 0.311 | 0.744 |
| **HEL × RYT** | Girdle | Female | 7 | 1 | 52.8 | No | 1.08e-04 | 0.024 | 0.056 | 0.263 | 0.815 |
| **HEL × RYT** | Spine | Male | 7 | 1 | 60.1 | No | 1.27e-07 | <0.001 | 0.103 | 0.285 | 0.892 |
| **HEL × RYT** | Spine | Female | 7 | 1 | 60.1 | No | 2.97e-08 | <0.001 | 0.124 | 0.311 | 0.937 |
| **HEL × RYT** | Girdle | male | 7 | 1 | 60.1 | No | 1.19e-06 | <0.001 | 0.094 | 0.311 | 0.995 |
| **HEL × RYT** | Girdle | female | 7 | 1 | 60.1 | No | 5.68e-07 | <0.001 | 0.103 | 0.263 | 1.025 |
| **HEL × RYT** | Spine | male | 7 | 1 | 70.7 | No | 1.48e-12 | <0.001 | 0.16 | 0.285 | 1.087 |
| **HEL × RYT** | Spine | female | 7 | 1 | 70.7 | No | 1.77e-13 | <0.001 | 0.212 | 0.311 | 1.141 |
| **HEL × RYT** | Spine | dom | 7 | 1 | 70.7 | No | 1.08e-07 | <0.001 | 0.086 | 0.213 | 0.814 |
| **HEL × RYT** | Girdle | male | 7 | 1 | 70.7 | No | 3.73e-11 | <0.001 | 0.145 | 0.311 | 1.249 |
| **HEL × RYT** | Girdle | female | 7 | 1 | 70.7 | No | 4.76e-12 | <0.001 | 0.191 | 0.263 | 1.316 |
| **HEL × RYT** | Girdle | dom | 7 | 1 | 70.7 | No | 1.35e-04 | 0.028 | 0.046 | 0.154 | 0.714 |
| **HEL × RYT** | Spine | male | 7 | 1 | 77.3 | No | 3.83e-24 | <0.001 | 0.201 | 0.285 | 1.383 |
| **HEL × RYT** | Spine | female | 7 | 1 | 77.3 | No | 1.96e-22 | <0.001 | 0.267 | 0.311 | 1.315 |
| **HEL × RYT** | Spine | dom | 7 | 1 | 77.3 | No | 2.53e-17 | <0.001 | 0.15 | 0.213 | 1.146 |
| **HEL × RYT** | Girdle | male | 7 | 1 | 77.3 | No | 1.51e-23 | <0.001 | 0.221 | 0.311 | 1.705 |
| **HEL × RYT** | Girdle | female | 7 | 1 | 77.3 | No | 6.99e-19 | <0.001 | 0.238 | 0.263 | 1.478 |
| **HEL × RYT** | Girdle | dom | 7 | 1 | 77.3 | No | 7.39e-12 | <0.001 | 0.103 | 0.154 | 1.134 |
| **HEL × RYT** | Spine | male | 7 | 1 | 83.7 | No | 6.13e-58 | <0.001 | 0.299 | 0.285 | 1.654 |
| **HEL × RYT** | Spine | female | 7 | 1 | 83.7 | No | 8.67e-55 | <0.001 | 0.312 | 0.311 | 1.573 |
| **HEL × RYT** | Spine | dom | 7 | 1 | 83.7 | No | 5.25e-48 | <0.001 | 0.215 | 0.213 | 1.432 |
| **HEL × RYT** | Girdle | male | 7 | 1 | 83.7 | No | 1.47e-48 | <0.001 | 0.311 | 0.311 | 2.041 |
| **HEL × RYT** | Girdle | female | 7 | 1 | 83.7 | No | 2.19e-38 | <0.001 | 0.277 | 0.263 | 1.711 |
| **HEL × RYT** | Girdle | dom | 7 | 1 | 83.7 | No | 3.59e-30 | <0.001 | 0.156 | 0.154 | 1.463 |
| **HEL × BYN** | Spine | male | 15 | 2 | 26.5 | No | 4.39e-06 | 0.001 | 0.07 | 0.095 | 0.466 |
| **HEL × BYN** | Spine | male | 15 | 2 | 33.3 | No | 1.12e-06 | <0.001 | 0.075 | 0.095 | 0.49 |
| **HEL × BYN** | Spine | male | 15 | 2 | 42.4 | No | 1.81e-07 | <0.001 | 0.09 | 0.095 | 0.521 |

MOLECULAR ECOLOGY

**Supplementary Table 3 | Detailed QTL results. Part 2/3.**

| Cross | Trait | Coding | LG | QTL | Pos | Std. | *P* | $P_{COR}$ | PVE | $PVE_{TOT}$ | β |
|---|---|---|---|---|---|---|---|---|---|---|---|
| **HEL × BYN** | Spine | male | 15 | 2 | 19.4 | No | 6.12e-06 | 0.001 | 0.061 | 0.095 | 0.465 |
| **HEL × BYN** | Spine | male | 15 | 2 | 26.5 | No | 4.39e-06 | 0.001 | 0.07 | 0.095 | 0.466 |
| **HEL × BYN** | Spine | male | 15 | 2 | 33.3 | No | 1.12e-06 | <0.001 | 0.075 | 0.095 | 0.49 |
| **HEL × BYN** | Spine | male | 15 | 2 | 42.4 | No | 1.81e-07 | <0.001 | 0.09 | 0.095 | 0.521 |
| **HEL × BYN** | Spine | male | 15 | 2 | 49.1 | No | 1.08e-07 | <0.001 | 0.088 | 0.095 | 0.529 |
| **HEL × BYN** | Spine | male | 15 | 2 | 56.1 | No | 9.00e-08 | <0.001 | 0.085 | 0.095 | 0.534 |
| **HEL × BYN** | Spine | male | 15 | 2 | 65.1 | No | 5.17e-08 | <0.001 | 0.088 | 0.095 | 0.54 |
| **HEL × BYN** | Spine | male | 15 | 2 | 77.5 | No | 5.05e-07 | <0.001 | 0.077 | 0.095 | 0.498 |
| **HEL × BYN** | Spine | male | 16 | 3 | 14.5 | No | 1.16e-04 | 0.01 | 0.054 | 0.086 | 0.397 |
| **HEL × BYN** | Spine | male | 16 | 3 | 19.9 | No | 1.21e-06 | <0.001 | 0.078 | 0.086 | 0.493 |
| **HEL × BYN** | Spine | male | 16 | 3 | 26.8 | No | 8.43e-07 | <0.001 | 0.082 | 0.086 | 0.493 |
| **HEL × BYN** | Spine | male | 16 | 3 | 35.8 | No | 2.59e-06 | 0.001 | 0.081 | 0.086 | 0.462 |
| **HEL × BYN** | Spine | male | 16 | 3 | 44 | No | 1.76e-06 | <0.001 | 0.067 | 0.086 | 0.47 |
| **HEL × BYN** | Spine | male | 16 | 3 | 56.5 | No | 9.25e-07 | <0.001 | 0.073 | 0.086 | 0.482 |
| **HEL × BYN** | Spine | male | 16 | 3 | 67.4 | No | 1.29e-06 | <0.001 | 0.071 | 0.086 | 0.475 |
| **HEL × BYN** | Spine | male | 16 | 3 | 82.2 | No | 1.72e-06 | <0.001 | 0.07 | 0.086 | 0.473 |
| **HEL × BYN** | Spine | male | 16 | 3 | 92.9 | No | 1.32e-06 | <0.001 | 0.07 | 0.086 | 0.48 |
| **HEL × BYN** | Spine | male | 16 | 3 | 102.2 | No | 4.18e-06 | 0.001 | 0.068 | 0.086 | 0.462 |
| **HEL × BYN** | Spine | male | 21 | 4 | 36.9 | No | 3.36e-04 | 0.031 | 0.037 | 0.037 | 0.36 |
| **HEL × BYN** | Spine | female | 6 | 5 | 41.2 | No | 1.37e-04 | 0.018 | 0.046 | 0.07 | 0.395 |
| **HEL × BYN** | Spine | female | 6 | 5 | 52 | No | 9.67e-06 | <0.001 | 0.061 | 0.07 | 0.458 |
| **HEL × BYN** | Spine | female | 6 | 5 | 59.4 | No | 1.08e-05 | <0.001 | 0.058 | 0.07 | 0.448 |
| **HEL × BYN** | Spine | female | 6 | 5 | 68.9 | No | 1.74e-06 | <0.001 | 0.067 | 0.07 | 0.5 |
| **HEL × BYN** | Girdle | male | 14 | 6 | 3.2 | No | 2.49e-04 | 0.024 | 0.038 | 0.039 | 0.306 |
| **HEL × BYN** | Girdle | male | 14 | 6 | 13.9 | No | 2.39e-04 | 0.023 | 0.05 | 0.039 | 0.307 |
| **HEL × PYÖ** | Spine | male | 9 | 7 | 3.4 | No | 8.96e-11 | <0.001 | 0.116 | 0.108 | 0.352 |
| **HEL × PYÖ** | Spine | male | 9 | 7 | 10.3 | No | 4.28e-10 | <0.001 | 0.101 | 0.108 | 0.348 |
| **HEL × PYÖ** | Spine | male | 9 | 7 | 19.2 | No | 1.70e-08 | <0.001 | 0.084 | 0.108 | 0.323 |
| **HEL × PYÖ** | Spine | male | 9 | 7 | 27.6 | No | 1.15e-08 | <0.001 | 0.09 | 0.108 | 0.33 |
| **HEL × PYÖ** | Spine | male | 9 | 7 | 44.7 | No | 1.68e-06 | <0.001 | 0.066 | 0.108 | 0.271 |
| **HEL × PYÖ** | Spine | male | 9 | 7 | 53.2 | No | 1.95e-04 | 0.024 | 0.042 | 0.108 | 0.209 |
| **HEL × PYÖ** | Spine | male | 9 | 7 | 64.8 | No | 3.97e-04 | 0.048 | 0.037 | 0.108 | 0.203 |

**Supplementary Table 3 | Detailed QTL results. Part 3/3.**

| Cross | Trait | Coding | LG | QTL | Pos | Std. | *P* | $P_{COR}$ | PVE | $PVE_{TOT}$ | β |
|---|---|---|---|---|---|---|---|---|---|---|---|
| **HEL × PYÖ** | Girdle | male | 19 | 8 | 1 | No | 1.56e-05 | 0.001 | 0.056 | 0.056 | 0.325 |
| **HEL × PYÖ** | Girdle | female | 4 | 9 | 0.2 | No | 1.32e-04 | 0.02 | 0.044 | 0.045 | 0.284 |
| **HEL × PYÖ** | Girdle | female | 4 | 9 | 2.5 | No | 1.60e-04 | 0.027 | 0.051 | 0.045 | 0.281 |
| **HEL × PYÖ** | Girdle | female | 4 | 9 | 8.5 | No | 1.55e-04 | 0.026 | 0.045 | 0.045 | 0.284 |
| **HEL × PYÖ** | Girdle | female | 4 | 9 | 13.9 | No | 7.11e-05 | 0.01 | 0.048 | 0.045 | 0.301 |
| **HEL × PYÖ** | Girdle | female | 4 | 9 | 19.1 | No | 1.28e-04 | 0.019 | 0.044 | 0.045 | 0.294 |
| **HEL × BYN** | Spine | male | 16 | 10 | 8.9 | Yes | 4.31e-04 | 0.048 | 0.037 | 0.098 | 0.007 |
| **HEL × BYN** | Spine | female | 16 | 10 | 26.8 | Yes | 2.53e-04 | 0.03 | 0.05 | 0.047 | 0.007 |
| **HEL × BYN** | Spine | female | 16 | 10 | 35.8 | Yes | 3.32e-04 | 0.044 | 0.04 | 0.047 | 0.007 |
| **HEL × BYN** | Girdle | female | 1 | 11 | 9.8 | Yes | 2.13e-05 | 0.002 | 0.048 | 0.077 | 0.005 |
| **HEL × BYN** | Girdle | male | 1 | 11 | 17.3 | Yes | 1.41e-04 | 0.016 | 0.026 | 0.026 | 0.004 |
| **HEL × BYN** | Girdle | female | 1 | 11 | 17.3 | Yes | 3.42e-06 | 0.002 | 0.052 | 0.077 | 0.005 |
| **HEL × BYN** | Girdle | male | 1 | 11 | 32.2 | Yes | 1.07e-04 | 0.014 | 0.023 | 0.026 | 0.004 |
| **HEL × BYN** | Girdle | female | 1 | 11 | 32.2 | Yes | 7.15e-08 | <0.001 | 0.077 | 0.077 | 0.006 |
| **HEL × BYN** | Girdle | male | 1 | 11 | 42.8 | Yes | 1.66e-05 | 0.001 | 0.028 | 0.026 | 0.005 |
| **HEL × BYN** | Girdle | female | 1 | 11 | 42.8 | Yes | 1.57e-07 | 0.001 | 0.062 | 0.077 | 0.006 |
| **HEL × BYN** | Girdle | male | 14 | 12 | 22.4 | Yes | 4.97e-05 | 0.006 | 0.041 | 0.041 | 0.004 |
| **HEL × BYN** | Girdle | male | 14 | 12 | 31.7 | Yes | 1.05e-04 | 0.013 | 0.041 | 0.041 | 0.004 |
| **HEL × BYN** | Girdle | male | 14 | 12 | 41.3 | Yes | 1.10e-04 | 0.014 | 0.041 | 0.041 | 0.004 |
| **HEL × PYÖ** | Spine | male | 1 | 13 | 32.2 | Yes | 4.75e-06 | <0.001 | 0.066 | 0.075 | 0.005 |
| **HEL × PYÖ** | Spine | male | 1 | 13 | 42.8 | Yes | 1.75e-06 | <0.001 | 0.078 | 0.075 | 0.005 |
| **HEL × PYÖ** | Spine | male | 1 | 13 | 57 | Yes | 2.30e-05 | 0.006 | 0.063 | 0.075 | 0.004 |
| **HEL × PYÖ** | Girdle | male | 1 | 13 | 57 | Yes | 9.69e-05 | 0.016 | 0.041 | 0.044 | 0.005 |
| **HEL × PYÖ** | Spine | male | 1 | 13 | 69.6 | Yes | 4.95e-05 | 0.008 | 0.058 | 0.075 | 0.004 |
| **HEL × PYÖ** | Girdle | male | 1 | 13 | 69.6 | Yes | 1.17e-04 | 0.018 | 0.046 | 0.044 | 0.005 |
| **HEL × PYÖ** | Spine | male | 1 | 13 | 79.2 | Yes | 1.12e-04 | 0.015 | 0.049 | 0.075 | 0.004 |

MOLECULAR ECOLOGY

**Supplementary Table 4 | Summary of candidate genes.** "Start" and "Stop" refer to the first and the last nucleotide positions in the nine-spined stickleback genome.

| Gene targetted | Gene found | Linkage group | Start | Stop |
|---|---|---|---|---|
| *Hoxd9[1] | Hoxd9b | LG6 | 1485512 | 1503619 |
| Hoxc9[1] | Na | na | na | na |
| Hoxb9[1,2] | Hoxb9a | LG11 | 14381929 | 14388893 |
| Hoxc10[1,2] | Na | na | na | na |
| Hoxc62 | Na | na | na | na |
| Wnt8c[1,2] | Na | na | na | na |
| Tbx4[1,2] | Tbx4 | LG12 | 10529939 | 10544947 |
| Tbx5[1,2] | Tbx5a | LG14 | 9729231 | 9742960 |
| Shh[1,2] | Shhb | LG12 | 8175862 | 8179502 |
| Shh[1,2] | Shhb | LG21 | 4618900 | 4626103 |
| Wnt3a[1,2] | Wnt3a | LG3 | 3158389 | 3168072 |
| *Fgf8[1,2] | Fgf8a | LG6 | 16496288 | 16500034 |
| *Fgf8[1,2] | Fgf8b | LG9 | 9040357 | 9043231 |
| Fgf10[1,2] | Fgf10 | LG8 | 17382172 | 17400365 |
| Fgf10[1,2] | Fgf10 | LG13 | 5433837 | 5439486 |
| Fgf10[1,2] | Fgf10 | LG14 | 8274848 | 8287244 |
| Fgf16[1] | Na | na | na | na |
| Fgf24[1] | Na | na | na | na |
| *Pel/Pitx1[1,2] | Pel/Pitx1 | LG7 | 7015000 | 17012000 |
| Pitx2[1,2] | Pitx2 | LG4 | 14810121 | 14819721 |
| *Pou1f1[3] | Pou1f1 | LG16 | 10088401 | 10094091 |
| *Hif1a[4] | Hif1a | LG15 | 7129489 | 7134581 |
| *Hif1a[4] | Hif1a | LG6 | 2836009 | 2847385 |
| Hif1a[4] | Hif1a | LG1 | 12028309 | 12049017 |

References: [1]Don *et al*. 2012; [2]Tanaka *et al*. 2005; [3]Kelberman *et al*. 2009; [4]Mudie *et al*. 201
Genes where no match was found are indicated with "*na*". Genes indicated with asterisk are in chromosomes with a significant QTL (Fig. 2, main document). References can be found in main document.

MOLECULAR ECOLOGY

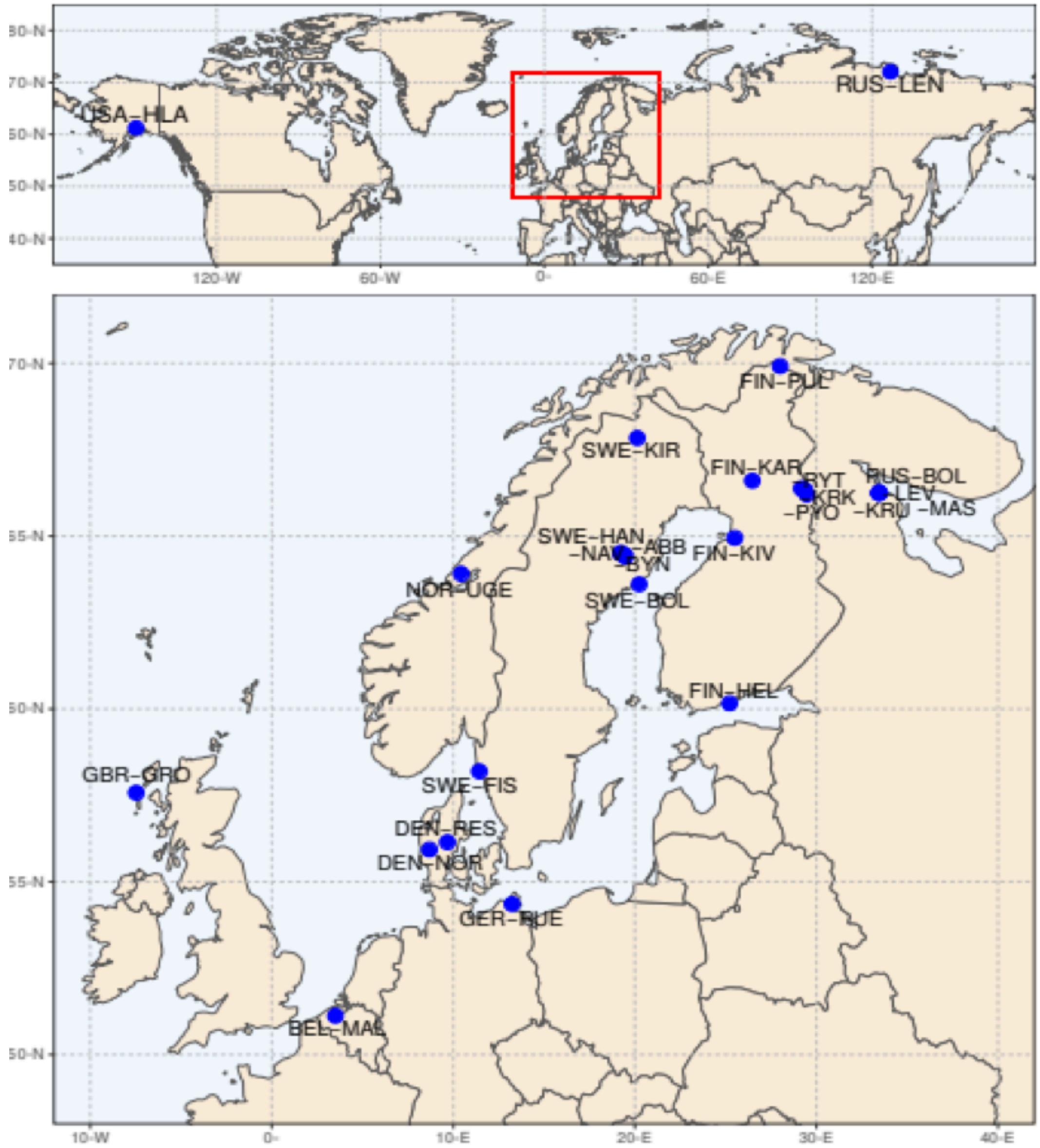


**Supplementary Figure 1 | Sample locations for *Pel*-deletion scan**

MOLECULAR ECOLOGY

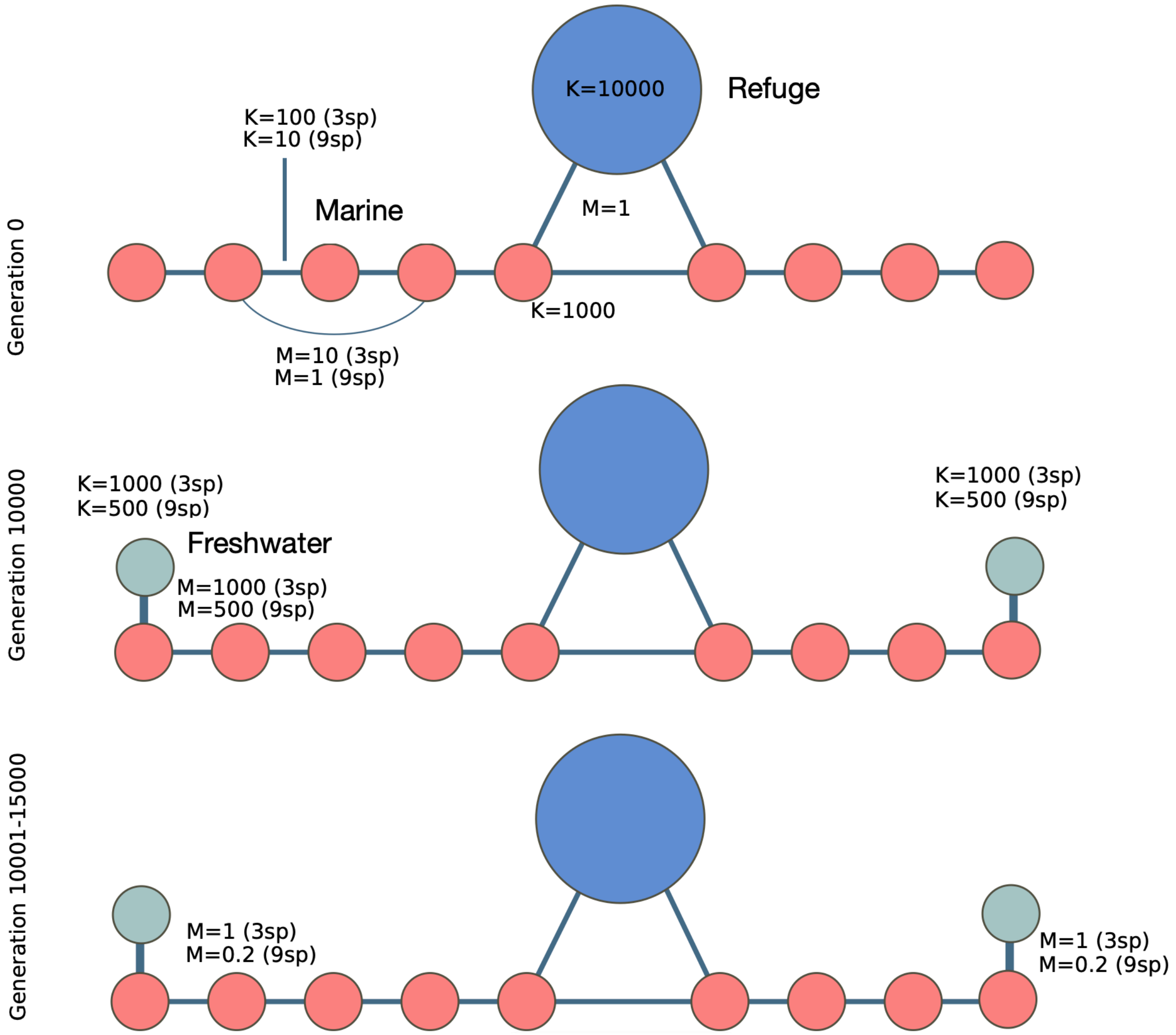


**Supplementary Figure 2 | Overview of proof of concept simulations.** Parameter settings are only shown when they change over generations (from top to bottom), and separately for three-spined (3sp) and nine-spined (9sp) sticklebacks. K is the carrying capacity and M is the number of migrants per generation.

MOLECULAR ECOLOGY

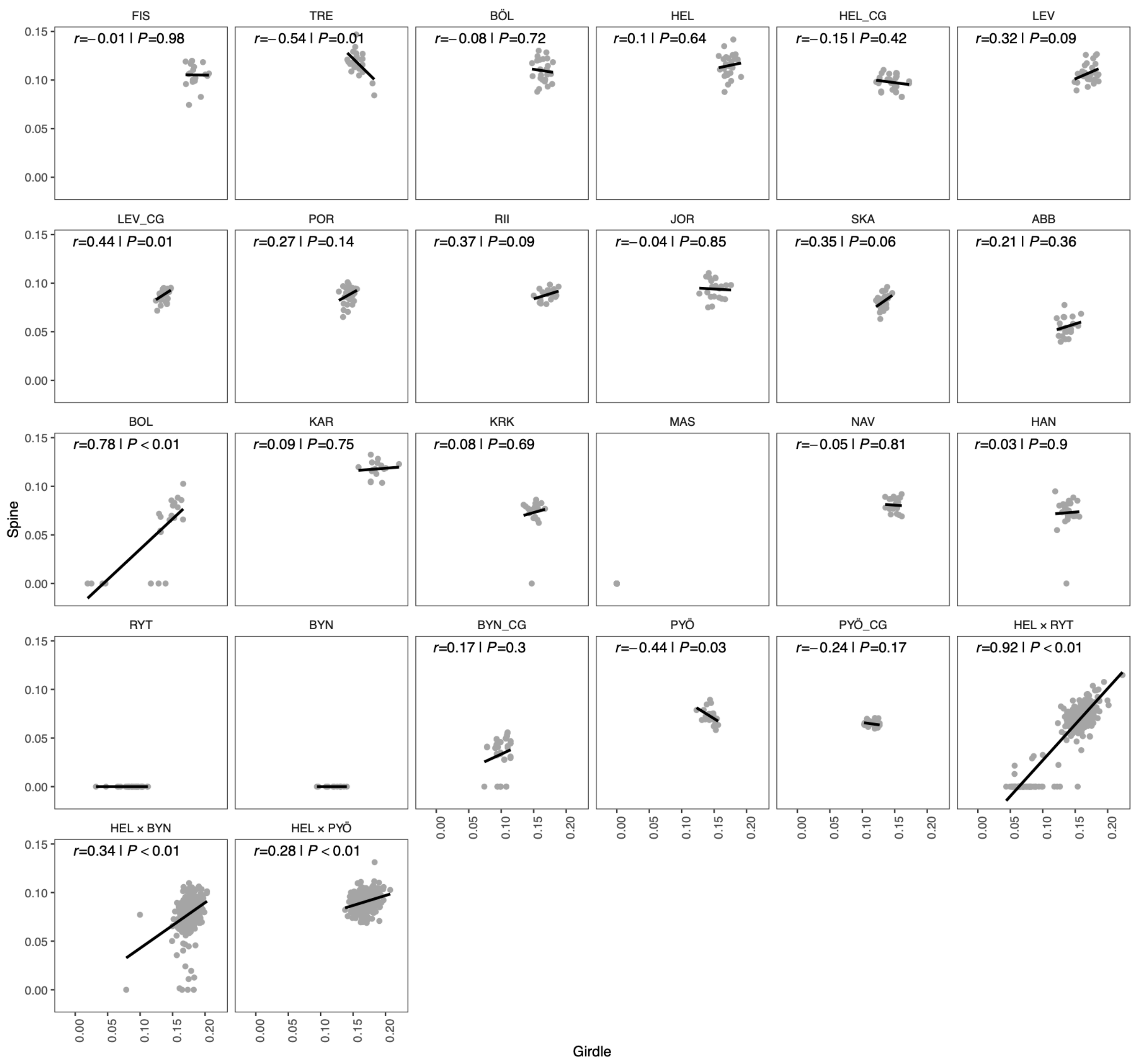


**Supplementary Figure 3 | Scatterplots between relative spine and girdle lengths.** Relative spine length is regressed against relative girdle length, with the correlation coefficients (*r*) and *P*-values indicated. Complements Figure 1 (main document) and Supplementary Table 1.

MOLECULAR ECOLOGY

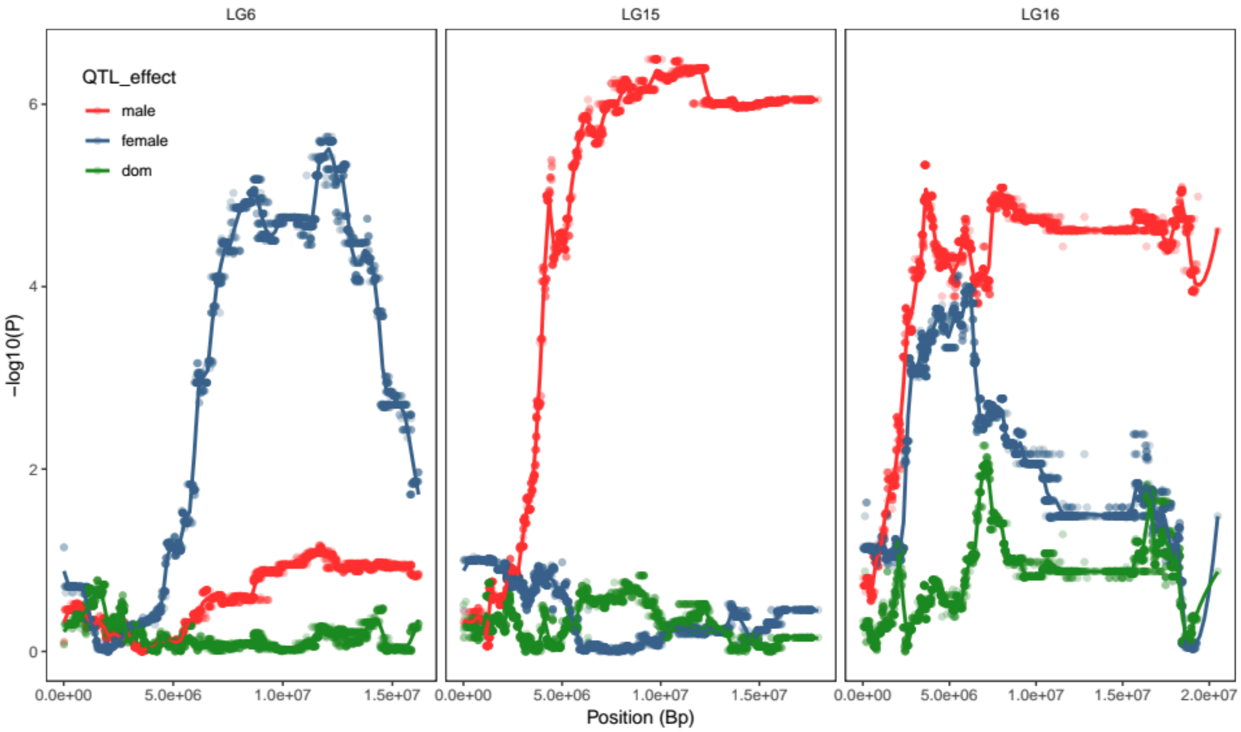


**Supplementary Figure 4 | Fine-mapping of pelvic spine length in LG6, LG15 and LG16.** Shown are nominal *P*-values from single-mapping four-way analyses independently for each SNP (no correction for multiple testing). A local regression (loess) has been fitted to the data (neighbourhood size = 0.05). The x-axis shows physical position in base pairs along each LG.

MOLECULAR ECOLOGY

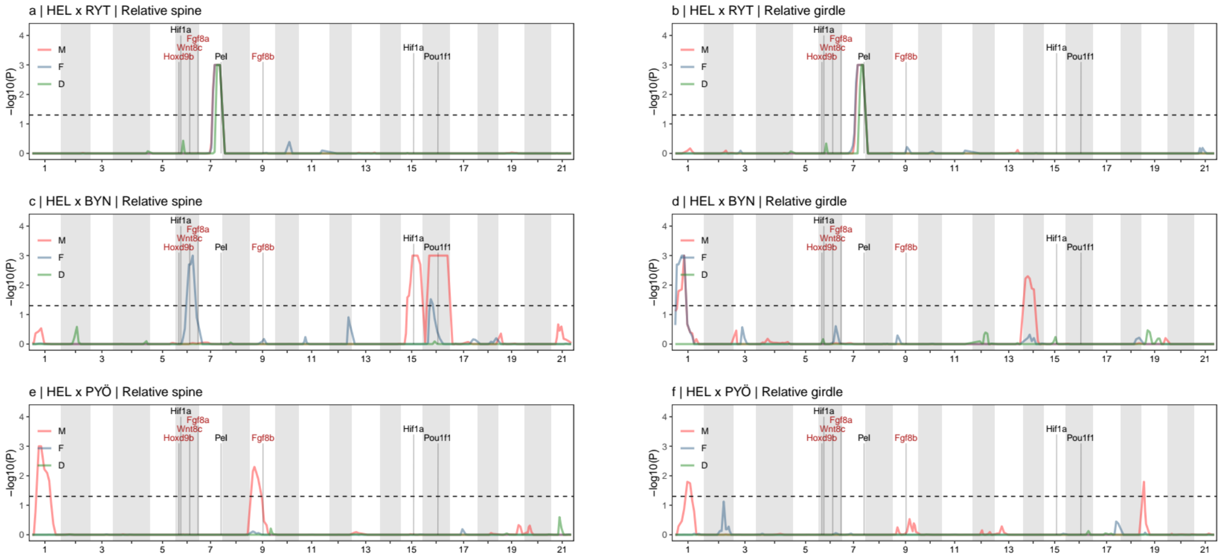


**Supplementary Figure 5 | Quantitative trait locus mapping of pelvic reduction, relative trait values.** Same as Figure 2, main text, except for relative trait values.

MOLECULAR ECOLOGY

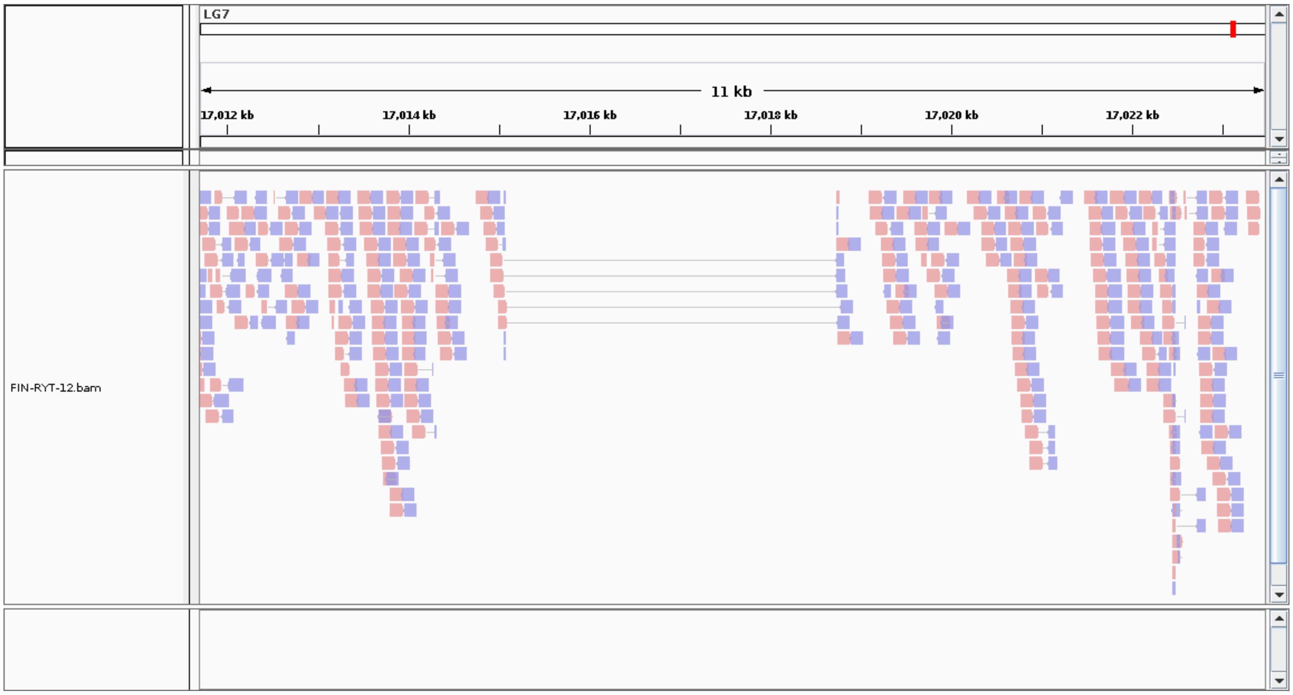


**Supplementary Figure 6 |** Alignment of split and clipped reads showing the exact location of the Pel-deletion (3,636 bp) found in Rytilampi pond.

MOLECULAR ECOLOGY

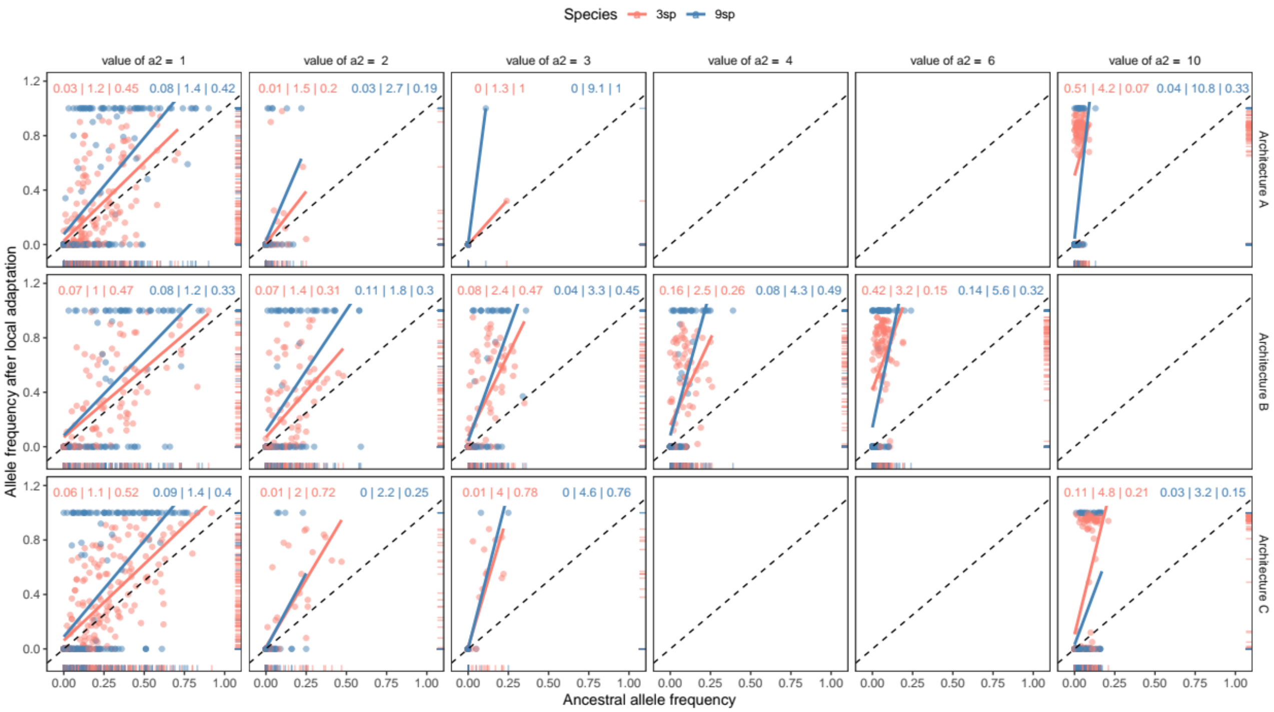


**Supplementary Figure 7 | Dependency of local adaptation on ancestral variation.** Shown are the ancestral allele frequencies of a2 (locally adapted to freshwater) in the marine population from which each freshwater population was founded at the time of colonisation, regressed against the frequency of a2 after local adaptation was complete (2k generations after colonisation). 3sp and 9sp refer to three-spined and nine-spined sticklebacks, respectively. Results are shown separately for QTL with different a2 effect sizes (columns) and architectures A, B and C (rows). Regression lines above 1 (shown as dashed line for reference) indicate that selection has increased a2 frequency in the freshwater population above that which was observed in the founding population. Higher slope and y-intercept indicates stronger effect of selection. The numbers in each panel show the y-intersection, regression slope and Pearson's squared correlation coefficient (separated by "|"), respectively, with colour indicating species. A strong correlation and the slope crossing the y-axis close to origo shows a strong dependence on ancestral variation for local adaptation and parallel evolution. Note that while e.g. for architecture A, the allele frequency of the QTL with a2 = 10 is close to fixed in most three-spined stickleback populations, the frequency of a2 for the QTL with effect size = 3 was almost never observed at high frequency in the freshwater population post local adaptation. Thus, interpreting the correlation for this figure is difficult. See also Supplementary File 7.

MOLECULAR ECOLOGY

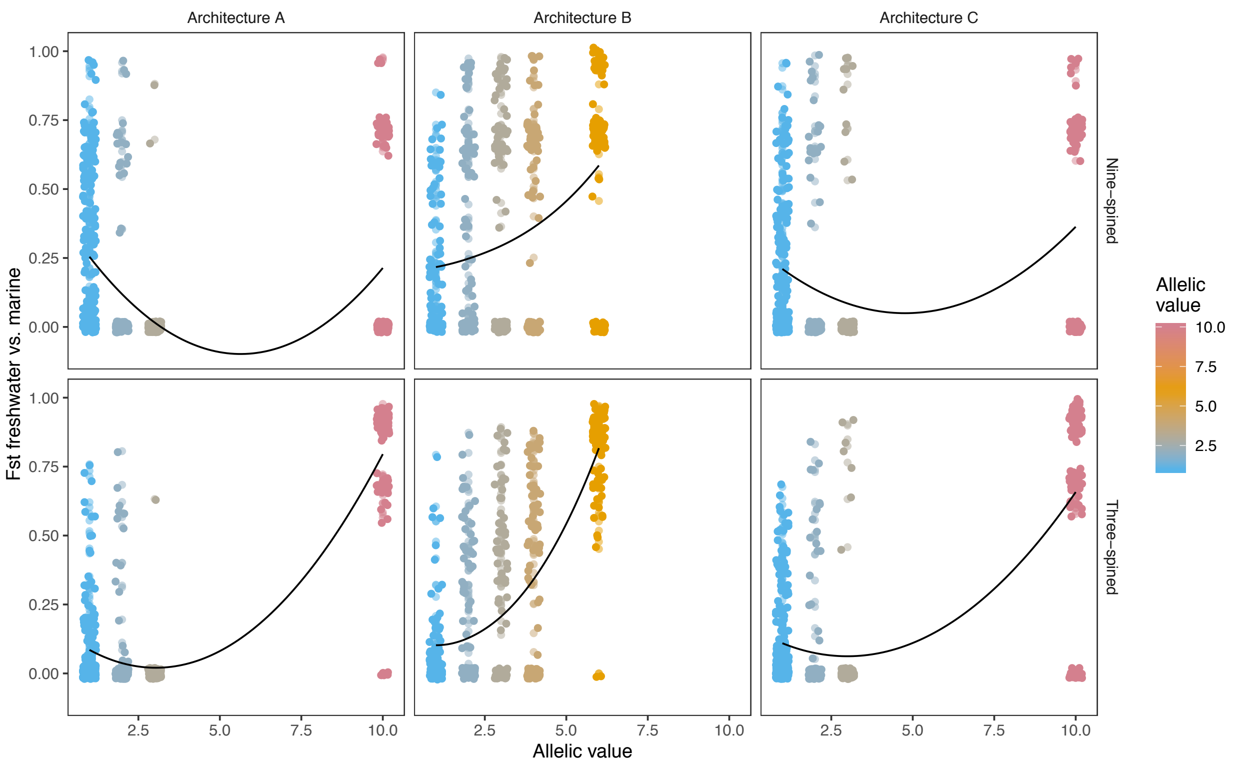


**Supplementary Figure 8 | QTL use in parallel evolution.** *F*-values between marine and freshwater populations as a function of QTL effect size (value of a2, the freshwater adapted allele). Colour scale is shown to aid comparison with Figure 5, main document. $F_{ST}$-values ~1 indicate the fixation of this allele in both of the two focal freshwater populations, and $F_{ST}$-values ~0.75 indicate fixation of the a2 allele in only one of them. Solid black line represents LOESS moving regression line. A single large effect QTL (as a homozygote) is enough to produce the optimal phenotype (a2 = 10, architecture A and C), and the medium effect QTL (most notably those with a2 = 2 or a2 = 3) were rarely involved in local adaptation. Because the smallest effect QTL (a2 = 1) did not have a large effect on the phenotype, their frequency was less affected by the large effect QTL. See also Supplementary File 7.
